# Hippocampal neuronal and astrocytic responses to noradrenaline and natural arousal

**DOI:** 10.64898/2026.01.16.699885

**Authors:** Sian N. Duss, Maria Wilhelm, Alina-Mariuca Marinescu, Runzhong Zhang, Fritjof Helmchen, Johannes Bohacek, Peter Rupprecht

## Abstract

The locus coeruleus (LC)-noradrenaline (NA) system is a central component of the brain’s response to arousal and stress. Yet how LC activity contributes to the cellular response profiles observed during natural arousal remains unclear. Here, we directly compared natural arousal with selective LC activation in mouse CA1, using physiologically titrated optogenetics, fiber photometry of NA and calcium signals, chronic two-photon imaging, and behavioral monitoring. While natural arousal robustly activated astrocytes, pyramidal cells, and interneurons, direct LC stimulation revealed a striking divergence in cellular response polarity and sensitivity. At levels of LC activation that correspond to arousal and moderate stress, we observed strong and reliable calcium responses in astrocytes, whereas pyramidal neurons and interneurons remained largely unaffected. Only at high-intensity LC stimulation did neurons exhibit a response, characterized by broad population-level inhibition of both pyramidal cells and interneurons, alongside the transient activation of an interneuron subpopulation that occupied distinct laminar positions in CA1. Thus, LC-driven NA release produces cell-specific effects in hippocampal CA1 that are distinct from and, at the population level for neurons even opposite to, cellular dynamics during natural arousal. Together, our results reveal a divergence in how astrocytes and neurons respond to LC-driven NA release and suggest that noradrenergic effects in the hippocampus at moderate levels of arousal and stress are predominantly mediated by astrocytes.

## Introduction

The arousal system in the brain is fundamental for our ability to flexibly recognize and respond to changes in the environment during rest, exploration, and stress. Readouts of arousal such as pupil dilations or unspecific motor activity have recently been shown to explain a large fraction of neuronal activity across the brain (McCormick et al., 2020; Musall et al., 2019; Stringer et al., 2019). These global signatures suggest that neuromodulatory systems play a key role in shaping arousal-related cellular activity. However, the specific contributions of individual neuromodulatory subsystems to these signals remain incompletely understood.

One of the central candidates is the locus coeruleus (LC), a small nucleus located in the pons and the principal source of noradrenaline (NA) in the brain. Through its highly divergent projections, LC broadcasts signals throughout virtually all regions of the central nervous system (Adér et al., 1980; Chandler et al., 2014). LC activity is tightly linked to changes in arousal, cognitive adaptation, and stress (Bouret and Sara, 2005; Devilbiss and Waterhouse, 2011; Vazey et al., 2018), and correlates closely with pupil dilations in both rodents and primates (Joshi et al., 2016; Privitera et al., 2020; Reimer et al., 2016). Despite this central role, it remains unclear how LC-mediated NA release acts at the cellular level in downstream regions and to what extent these direct effects contribute to the circuit dynamics responses observed during natural arousal.

The hippocampus (HC) is a stress-sensitive brain region that receives noradrenergic input exclusively from the LC. Its well-characterized circuitry and dense noradrenergic innervation make it an ideal system to study the cellular actions of LC-mediated NA-release (LC-NA). NA release strongly influences hippocampal function particularly during states of arousal or stress (Berridge and Waterhouse, 2003; Clewett et al., 2018; Giustino et al., 2020; Grueschow et al., 2020; Murchison et al., 2004; Roozendaal and McGaugh, 2011). Hippocampal activity is also strongly modulated during arousal and movement: pyramidal cells increase firing rates during locomotion in a speed-dependent manner (Czurkó et al., 1999; Fuhrmann et al., 2015; McNaughton et al., 1983), interneurons are robustly activated or, for a subset, inhibited during arousal (Arriaga and Han, 2017; Geiller et al., 2020; Lapray et al., 2012; Yang et al., 2025), and astrocytes exhibit slow centripetal activations upon movement and arousal (Rupprecht et al., 2024). These diverse, cell type-specific responses highlight that arousal profoundly shapes hippocampal dynamics. Yet it is not known to what degree these effects are driven directly by LC-NA release versus other arousal-related pathways.

Early work by Segal and Bloom in the 1970s showed that NA can strongly influence hippocampal activity, reporting NA-mediated inhibition in the hippocampus using electrical stimulation of the LC and iontophoretic application of NA in awake rats (Segal and Bloom, 1976, 1974a, 1974b). However, these pioneering experiments lacked the cell-type specificity required to disentangle the effects on the local circuit. More recent work, primarily in brain slices, has characterized the responses to NA of specific hippocampal cell types, describing an NA-mediated increase of excitability in pyramidal cells (Bacon et al., 2020; Church et al., 2019; Madison and Nicoll, 1986), an activation of interneurons (Bergles et al., 1996; Hillman et al., 2009; Madison and Nicoll, 1988), and a very prominent activation of astrocytes (Gao et al., 2016; Lefton et al., 2025). Yet most of these studies relied on *ex vivo* slices or exogenous NA application. Therefore, these findings demonstrate the strong influence of LC-NA on hippocampal circuits, but they do not reveal how physiological LC-NA release modulates identified cell types *in vivo*, nor to which extent these effects mimic arousal-driven activity in awake animals.

To clarify how LC-NA shapes hippocampal activity and to determine its contribution to the arousal response, we systematically investigated the effects of both natural arousal and selective LC-NA stimulation on hippocampal cell types in awake mice. We first assessed physiologically relevant NA levels during acute stress and arousal, and titrated optogenetic LC stimulation accordingly. Using a combination of LC stimulation, fiber photometry, chronic two-photon imaging in CA1, and behavioral monitoring, we then compared the response of pyramidal cells, interneurons, and astrocytes to natural arousal and selective LC-NA stimulation. This approach provides a comprehensive, cell type-specific analysis of LC-NA effects *in vivo* and enables a direct evaluation of the contribution of LC-NA to hippocampal arousal responses.

## Results

### Titration of LC stimulation to physiological NA levels

LC activity is known to scale with wakefulness and arousal, and reaches its highest firing rates during stress (Akaike, 1982; Aston-Jones and Bloom, 1981; Carter et al., 2010). To relate LC activity to physiologically relevant levels of NA release, we measured NA dynamics in hippocampal CA1 during both acute stress and LC stimulation using fiber photometry. We used virally induced expression of the optogenetic actuator ChrimsonR in the LC of DBH-iCre animals and of NA sensors (GRAB-NE2m in Figure 1; GRAB-NE1m and nLightG in Figure S1-4) in neurons of the HC (Figure 1A-C). We subjected animals to a short 10-s tail lift and a 3-s mild foot shock (Figure 1D). As previously reported (Feng et al., 2024, 2019), the NA level increased during the tail lift and gradually decayed afterwards (Figure 1E). The foot shock triggered NA signals with comparable peak amplitudes that, however, remained elevated throughout the time recorded (Figure 1F), in line with published work (Li et al., 2023). As additional control for potential movement artifacts, we repeated all experiments with expression of GFP instead of NA sensors but found no signal change in response to either stressor in these animals (Figure S1-1). To estimate the maximal range of NA release, animals were subjected to the cold forced swim paradigm, a stressor known to potently activate NA release (Ebner and Singewald, 2007; Privitera et al., 2024). This paradigm, compared to the other stressors, induced substantially higher peak NA levels that decayed only slowly (Figure S1-2). Finally, to confirm that the recorded NA signals originated from the LC, we optogenetically inhibited the LC during stress exposure using the chloride pump Jaws (Chuong et al., 2014). We found that the increase in NA levels in response to stressors was largely prevented by optogenetic inhibition of the LC (Figure S1-3). Together, these findings show that distinct stressors evoked graded, LC-dependent NA release in the hippocampus, providing a physiological reference for calibrating optogenetic LC stimulation.

**Figure 1.**
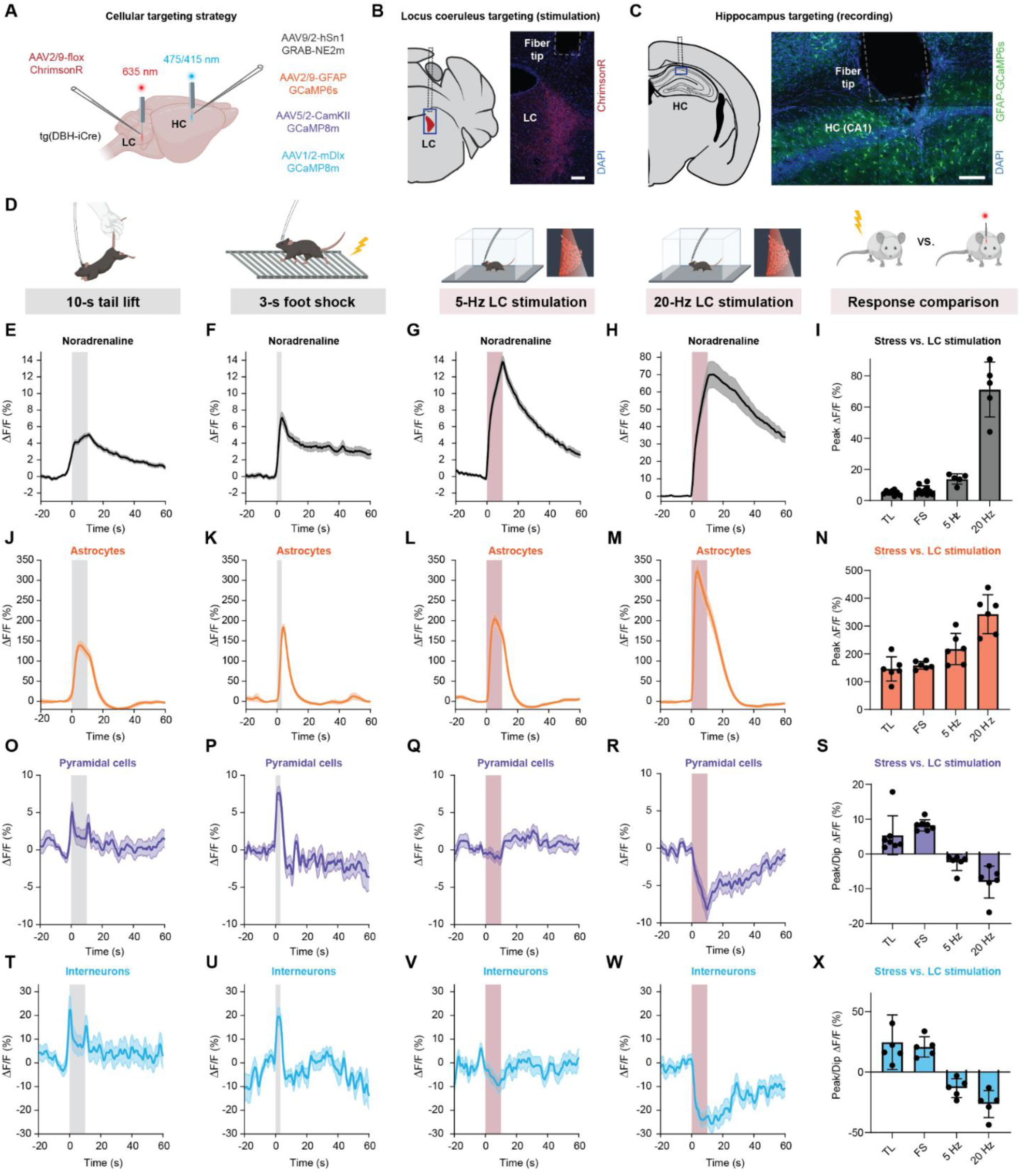
Optogenetic LC stimulation and stress trigger similar responses in astrocytes but not in excitatory or inhibitory neurons. A, Schematic representation of the viral strategy and experimental approach using fiber photometry recordings in hippocampus and optogenetic LC stimulation during behavior. **B,** Schematic and histology example of LC targeting using DBH-iCre transgenic mice together with injection of Cre-dependent AAV with ChrimsonR-tdTomato (red). Scale bar, 100 μm. **C,** Schematic and histology example of HC targeting using injection of AAV with, in this case, GFAP-GCaMP6s (green) to target astrocytes, visualized together with nuclear stain (blue, DAPI). Scale bar, 100 μm. **D,** Schematic representation of the experiments with corresponding signals below. **E-I,** NA release in response to (E) a 10-s tail lift (n = 10), (F) a 3-s foot shock (n = 10), (G) optogenetic 10-s 5-Hz LC stimulation (n = 5), and (H) 10-s 20-Hz stimulation (n = 5). Comparison of the peak ΔF/F in (I). **J-N,** Astrocytic Ca^2+^ in response to (J) a 10-s tail lift (n = 6), (K) a 3-s foot shock (n = 6), (L) optogenetic 10-s 5-Hz stimulation (n = 6), and (M) 10-s 20-Hz stimulation (n = 6). Comparison of the peak ΔF/F in (N). **O-S,** Pyramidal cell Ca^2+^ in response to (O) a 10-s tail lift (n = 7), (P) a 3-s foot shock (n = 7), (Q) 10-s 5-Hz stimulation (n = 6), and (R) 10-s 20-Hz stimulation (n = 6). Comparison of the peak ΔF/F in (S). **T-X,** Interneuron Ca^2+^ in response to (T) a 10-s tail lift (n = 6), (U) 3-s foot shock (n = 6), (V) 10-s 5-Hz stimulation (n = 5), and (W) 10-s 20-Hz stimulation (n = 5). Comparison of the peak ΔF/F in (X).

Next, we aimed to find LC stimulation strengths that approximate peak NA release during mild and intense stress exposure, respectively. First, we titrated LC-NA release under anesthesia and found that NA release increased with LC stimulation frequency from 1 Hz up to 40 Hz, where NA release plateaued at levels comparable to continuous LC stimulation (Figure S1-4B). As a proxy of mild stress, we applied LC stimulation at 5 Hz for 10 s (Figure 1G), a protocol previously linked to stress and anxiety (McCall et al., 2015; Privitera et al., 2024). To mimic peak NA levels observed under intense stress, we applied LC stimulation at 20 Hz (Figure 1H; Figure S1-4B). Together, these results establish 5-Hz and 20-Hz LC stimulation as approximations of LC-NA release observed during moderate and intense acute stress (Figure 1I).

### LC stimulation and natural stress elicit similar responses in astrocytes but distinct responses in both pyramidal cells and interneurons

To assess cellular responses in the HC, we next recorded from astrocytes, a cell type known for their sensitivity to stress and their abundant expression of adrenergic receptors (Hertz et al., 2010). We first used fiber photometry to record astrocytic Ca2+ responses (GFAP-GCaMP6s). When animals were subjected to stressors, we found a surge in astrocytic Ca2+ during the acute stress exposure that went back to baseline on a timescale of seconds (Figure 1J,K). Similarly, both LC stimulation paradigms (5 Hz and 20 Hz) triggered elevations in astrocytic Ca2+ (Figure 1L,M). Peak astrocyte Ca2+ signals (Figure 1N) followed a similar trend as peak NA levels (Figure 1I), albeit with a seemingly smaller dynamic range. This similarity of response profiles suggests a close relationship between stress, NA and astrocytic activation.

Next, we investigated neuronal responses to LC stimulation and natural stress. To this end, we expressed Ca2+ sensors in hippocampal pyramidal cells (CamKIIα-GCaMP8m) or GABAergic inhibitory neurons (mDlx-HBB-GCaMP8m) and used fiber photometry to record neuronal bulk responses during the same conditions as before. Both neuronal cell types displayed elevated Ca2+ signals during tail lift, predominantly at onset (Figure 1O,T), and during foot shock (Figure 1P,U). LC stimulation, in contrast, evoked a long-lasting decrease in Ca2+ that scaled with the intensity of the stimulation (Figure 1Q,R,V,W). Therefore, unlike NA levels and signals recorded from astrocytes, bulk Ca2+ signals from pyramidal cells and interneurons showed opposite responses to natural stress and optogenetically induced NA release (Figure 1S,X). Together, these fiber photometry results demonstrate a direct inhibitory effect of NA released from LC on neuronal activity, which, however, appears to be outweighed by other factors driving activity during natural stress. The increased neuronal activity during stress is therefore unlikely to be directly driven by release of NA upon LC activity. These analyses are, however, limited by the fact that fiber photometry reflects population-averaged signals and cannot resolve cell-specific contributions to neuronal or astrocytic activity, prompting us to examine single-cell activity patterns using two-photon calcium imaging.

### Cell-type specific two-photon imaging during LC stimulation and natural arousal

To investigate the functional diversity within populations of each cell type, we used two-photon calcium imaging in mice that were head-fixed on a treadmill (Figure 2A). Analogous to fiber photometry experiments, we expressed calcium sensors in astrocytes (n = 4 mice), pyramidal cells (n = 5) or interneurons (n = 4) and ChrimsonR in LC neurons. Then, we implanted an optical fiber targeting LC, and performed multi-plane calcium imaging of hippocampal CA1 through an implanted cannula, while repeatedly applying optogenetic LC stimulation of varying strength (5 Hz and 20 Hz for moderate and strong stimulation paradigms, or, in selected sessions, a broader range of stimulation intensities from 1 Hz to 40 Hz; additional 40 Hz stimulations were applied for pyramidal cells and interneurons, see Methods) (Figure 2B). Imaging was performed within a single imaging plane for pyramidal cells, across 3 planes for astrocytes, and across 4 planes for interneurons, with imaging planes spaced by 40 μm for multi-plane imaging. We therefore spanned a large fraction of *stratum oriens* together with *stratum pyramidale* for astrocytic imaging, and the entire *stratum oriens*, *stratum pyramidale* and a variable but relatively small part of *stratum radiatum* for interneuron imaging. Chronic calcium imaging was performed across multiple days (Figure 2C), enabling the tracking of individual cells and their activity during LC stimulation and natural arousal across days (Figure S2-1).

**Figure 2.**
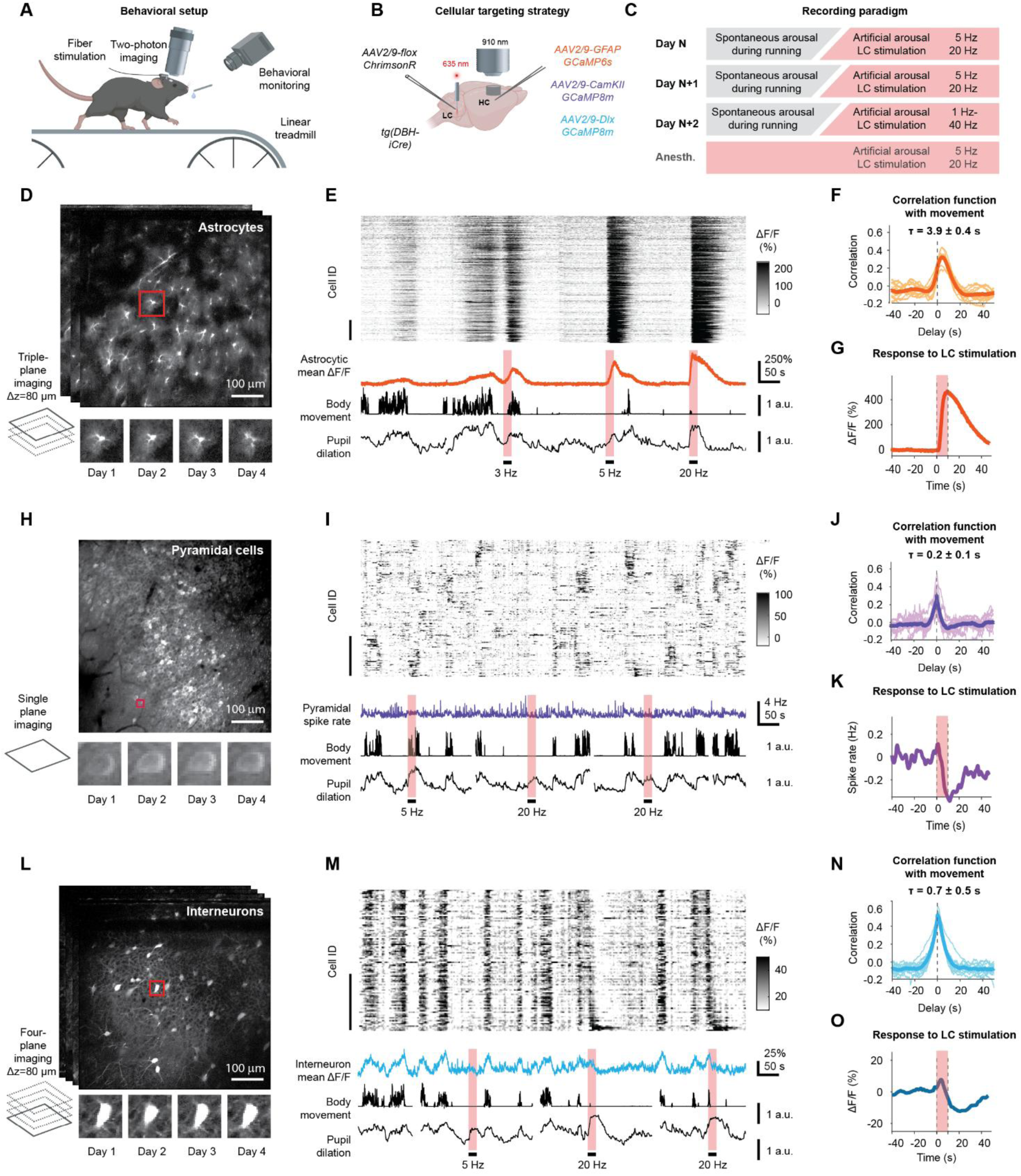
Chronic two-photon imaging during optogenetic LC stimulation and spontaneous behavior across neurons and astrocytes. **A**, Schematic representation of the two-photon recording setup. **B**, Viral strategy to stimulate the LC and record Ca2+ signals from either astrocytes (orange), excitatory neurons (purple) or inhibitory neurons (turquoise). **C**, Recording paradigms across the imaging days. **D**, *Top:* Representative FOV of triple-plane imaging in astrocytes. *Bottom:* Example cell tracked across recording days. **E**, Example recording from astrocytes. From top to bottom: sorted time traces of all cells identified in (D) (n = 310 cells; the vertical scale bar, here and in panels (I) and (M), represents 50 cells); average ΔF/F across cells; body movement and pupil diameter extracted from behavioral tracking video. LC stimulations are indicated with a red shaded bar (10 s stimulation at 3 Hz, 5 Hz or 20 Hz). **F**, Correlation function of body movement and astrocytic Ca^2+^ signals, averaged across all animals and sessions in dark color, individual sessions (n = 12) indicated in lighter color. The delay τ measures the average delay (median ± S.D. across sessions) of the calcium signal compared to body movement. **G**, Mean astrocyte response to 20 Hz LC stimulation, averaged across all stimulations (n = 32 stimulations) and animals (N = 4). **H**, *Top:* Representative FOV of single-plane imaging in excitatory neurons. *Bottom:* Example cell tracked across recording days. **I**, Example recording from pyramidal cells. From top to bottom: rastermap sorting of time traces from selected cells (n = 170 out of 513 detected cells); spike rate estimate averaged across all detected cells; body movement and pupil diameter extracted from behavioral tracking video. LC stimulations are indicated with a red shaded bar (10 s stimulation at 5 Hz or 20 Hz). **J**, Correlation function of body movement and pyramidal cell Ca^2+^ signals, averaged across all animals and sessions (n = 16) in dark color, individual sessions indicated in lighter color. **K**, Mean pyramidal cell response to 20 Hz LC stimulation, averaged across all stimulations (n = 36 stimulations) and animals (N = 5). **L**, *Top:* Representative images of the four-plane imaging in interneurons. *Bottom:* Example cell tracked across recording days. **M**, Example recording from interneurons. From top to bottom: sorted time traces of all cells identified in (L) (n = 124 cells); average ΔF/F across cells; body movement and pupil diameter extracted from behavioral tracking video. LC stimulations are indicated with a red shaded bar (10 s stimulation at 5 Hz or 20 Hz). **N**, Correlation function of body movement and interneuronal Ca^2+^ signals, averaged across all animals and sessions (n = 14) in dark color, individual sessions indicated in lighter color. **O**, Mean interneuron response to 20 Hz LC stimulation, averaged across all stimulations (n = 39 stimulations) and animals (N = 4).

To compare the effects of LC stimulation and natural arousal, we first needed a reliable proxy for endogenous arousal. Earlier work has shown that both locomotion and pupil diameter can be used as proxies for arousal state (McGinley et al., 2015; Shimaoka et al., 2018; Stringer et al., 2019). In our head-fixed experiments, mice exhibited spontaneous behaviors that included episodes of both low arousal (resting) and high arousal (accompanied by running on the treadmill), monitored by camera recordings of body movements and pupil diameter. We and others have previously shown for head-fixed mice that face or body movement, more so than locomotion alone, is highly predictive of astrocytic activity and arousal (Collins et al., 2023; Rupprecht et al., 2024), which we confirm here by predicting pupil diameter with body movement (Figure S2-2). In addition, pupil recordings, more so than recordings of body movement, were prone to noise due to pupil dilations upon optogenetic LC stimulation (Figure S2-3A) or eye movements. Interestingly, LC stimulation not only resulted in pupil dilations but also, for stronger LC stimulation, sometimes resulted in reduced body movement (Figure S2-3B), consistent with previous reports of behavioral arrest during strong and prolonged LC stimulation (Carter et al., 2010). We therefore decided to use body movement as our main proxy of endogenous arousal. Together, our experimental paradigm enabled us to monitor arousal while recording cellular activity (see also Supplementary Movies 1, 2 and 3).

We find that astrocytic calcium signals were largely synchronized across the field of view (FOV) during body movement and upon optogenetic LC stimulation (Figure 2D,E; Figure S2-4A,B), consistent with previous findings in cortex and hippocampus (Ding et al., 2013; Paukert et al., 2014; Rupprecht et al., 2024). Astrocytic responses to body movement as well as responses to optogenetic LC stimulation during wakefulness and anesthesia exhibited a strictly positive modulation when averaged across cells within a session (Figure 2F,G; Figure S2-5), but also on the level of individual astrocytes (Figure S2-5A,B). These findings are consistent with our bulk fiber photometry recordings (Figure 1, Figure S2-6) and reflect the well-described property of astrocytic populations as a slow and synchronized tracker of past arousal or LC activity (Fedotova et al., 2023; Rupprecht et al., 2024).

Calcium imaging of neurons in the densely packed pyramidal layer of CA1 displayed cellular activity that evolved on a much faster timescale and in a much less synchronized manner, without any obvious relationship between behavioral proxies of arousal and pyramidal cell activity (Figure 2H,I). After denoising by spike rate inference of the cellular calcium imaging data (Rupprecht et al., 2021) and averaging across neurons, we found an overall positive modulation of population activity with movement (Figure 2J), consistent with fiber photometry (Figure 1; Figure S2-6) and previous work (Rupprecht et al., 2024). On the single-neuron level, a minority of pyramidal cells was inhibited during movement (Figure S2-4A). In contrast, and consistent with fiber photometry recordings (Figure 1), we observed a marked decrease of neuronal activity in response to strong stimulation of LC (Figure 2K).

Calcium signals in interneurons were more synchronized across cells compared to pyramidal cells and clearly modulated by body movement as behavioral proxies for arousal (Figure 2L,M,N, S2-4A,B; Figure S2-6). Notably, and in contrast to astrocytic responses, the similarity of responses within imaging planes was higher than the similarity of responses across imaging planes, demonstrating a laminar organization of response characteristics in hippocampal CA1 for interneurons but not astrocytes (Figure S2-4C,D). Interestingly, interneurons exhibited, on average, a slow decrease of activity upon LC stimulation, consistent with fiber photometry, but this decrease was preceded by a transient increase in overall activity (Figure 2O; Figure S2-4A). As for pyramidal cells, these effects scaled with stimulation intensity.

Thus, two-photon imaging enables detailed analyses of cellular response characteristics across cell types and highlights the different response characteristics, temporal scales and synchronization regimes of these three major cell types under otherwise identical recording conditions.

### Astrocytes exhibit diverse but stable cell-specific sensitivity to LC stimulation

It is well-established that astrocytes can be activated by NA released from LC axons (Ding et al., 2013; Paukert et al., 2014). In agreement with this, we observe that virtually all astrocytes were activated by LC stimulation, albeit to varying degrees (Figure 3A). We wondered whether this degree of activation is a property that is specific to each astrocyte or whether it depends on other factors (e.g., the current state of the cell). We found that the current state of each cell (pre-stimulus ΔF/F level) explained only a small fraction of the variance of responses (Figure 3B; correlation *r* = 0.17). In contrast, the identity of a cell explained a much larger fraction of the variance, both within sessions (Figure 3C,D, *r* = 0.56, pooled across 4 mice) as well as across days (Figure 3E, *r* = 0.61 ± 0.12, mean ± S.D. across 4 mice). This provides direct evidence for a consistent and cell-specific responsiveness to LC stimulation. In parallel, we observed that the modulation of the activity patterns of individual astrocytes by body movement (as a proxy for arousal) was also conserved across days for individual astrocytes (Figure 3F, *r* = 0.56 ± 0.29). These results complement the recently described morphological and transcriptomic diversity of astrocytic cell types (Batiuk et al., 2020; Endo et al., 2022; Viana et al., 2023) from a functional perspective, based on each cell’s responsiveness to natural arousal and NA release.

**Figure 3.**
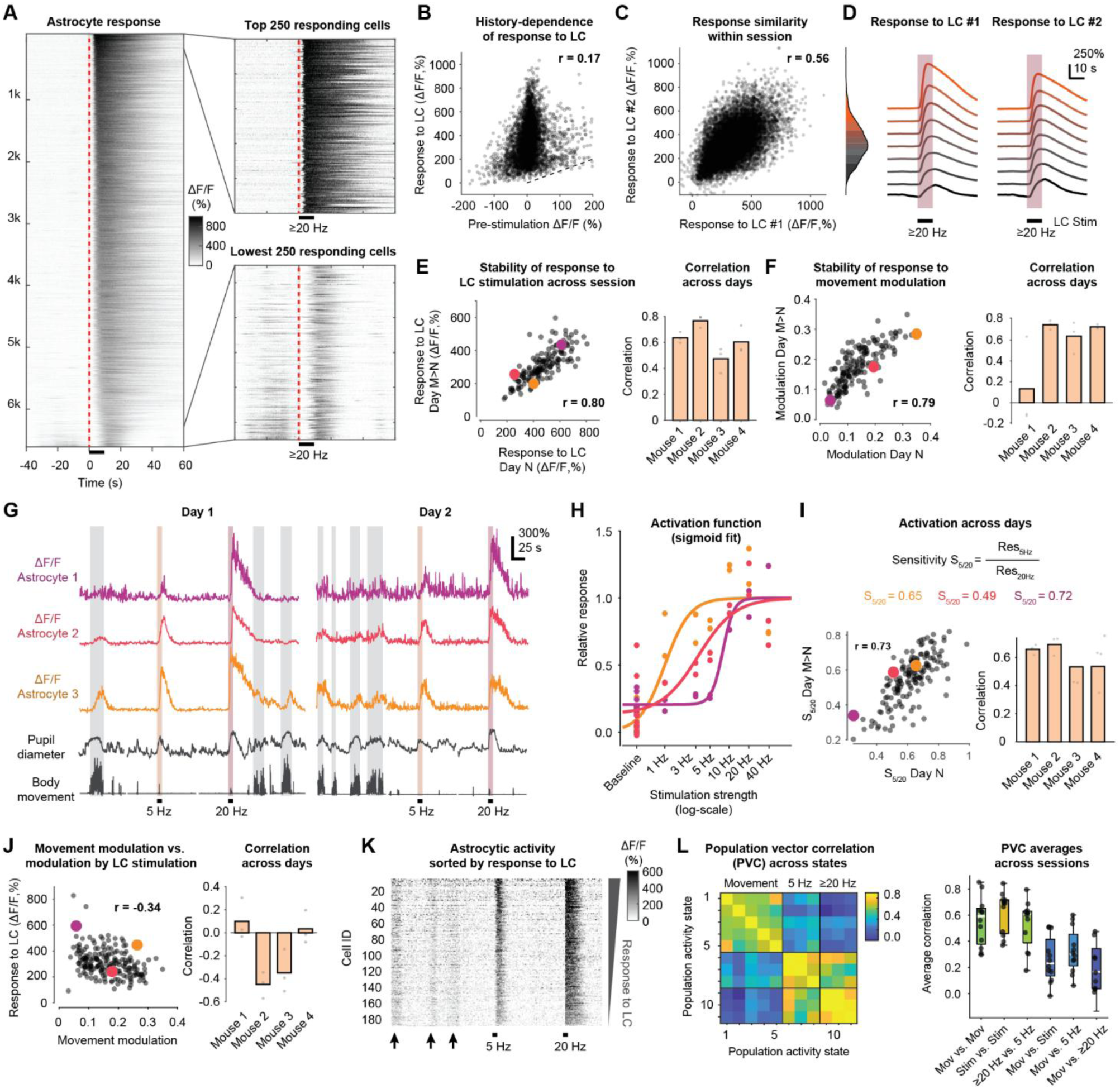
Astrocyte subpopulations exhibit stable, cell-specific responses to LC stimulation that diverge from activity during natural arousal. **A**, All astrocytic responses to ≥20 Hz LC stimulation, sorted by the ΔF/F change during the 10 s stimulation window compared to the 20 s window before stimulation, with subselection of the 250 most and least responsive cells. Data pooled from 12 sessions across 4 mice. **B**, Time-averaged response to a strong stimulation as shown in panel (A) as a function of pre-stimulus ΔF/F (average across 20 s before stimulation). **C**, *Left:* Response to two strong LC stimulations within a single session (stimulation 1: x-axis, stimulation 2: y-axis) to investigate within-session stability. *Right:* Marginal distribution of LC stimulation responses, colorcoded as in panel (D). **D**, Analysis of stability of astrocytic responses within sessions. *Left:* Cellular responses to stimulation #1, sorted by most to least responsive cells and plotted as the quantiles shown in panel (C, right). *Right:* Responses of the same cells to stimulation #2 within the same session, with the sorting maintained from stimulation #1. **E**, *Left:* Analysis of stability of astrocytic responses towards LC stimulation for an individual session pair (day N vs. day N+1). Here and in panels (H,I,J), only a single session pair is shown as scatter plot due to large differences between mean average response magnitudes across sessions and animals. *Right:* Correlation of responses across days for each animal and each session pair. Here and in the following panels, the colored dots correspond to the astrocytes shown in panel (G). **F**, *Left:* Analysis of stability of astrocytic activity locked to body movement across a session pair (day N vs. day N+1). *Right:* Correlation of responses across days for each animal and each session pair. **G**, Examples of distinct astrocytic response types. Episodes of natural arousal encompassed by body movement are highlighted with gray shading (manual annotation). Episodes of optogenetic stimulation of LC are highlighted with red shading. Simultaneously recorded body movement as a proxy for natural arousal is shown below. The same astrocytes are shown for day 1 (left) and day 2 (right). **H**, Sigmoid fit of activation in response to increasing LC stimulation frequency for the three selected astrocytes displayed in (G). Activation functions are shown normalized to the saturating response of each astrocyte. **I**, Response sensitivity of astrocytes to LC stimulation, defined as the ratio of responses to 5 Hz vs. 20 Hz stimulations of LC. *Left:* Example stability of response sensitivity across two imaging sessions. *Right:* Pearson correlation coefficient for each session pair for all animals. **J**, *Left:* Correlation of responses to LC stimulation and movement modulation. *Right:* Pearson correlation coefficient for all four animals. **K**, Example activity traces from one mouse, ΔF/F sorted by LC stimulation response (same segment as for Day 2 in F.). Black arrows indicate bouts of body movement. **L**, *Left: E*xample of population vector correlations for one recording session comparing movement episodes with instances of 5 Hz and 20 Hz LC stimulation. *Right:* Average Pearson correlation coefficient for each comparison. Each data point indicates an imaging session.

To investigate astrocytic response types in more depth, we closely inspected their response patterns to LC stimulations of varying intensity. We noticed that some astrocytes already responded strongly to stimulation of intermediate strength (5 Hz), while others were only activated upon strong LC stimulation (20 Hz) (Figure 3G). In a subset of experiments, we covered a large range of stimulus strengths (ranging from 1 Hz to 40 Hz), enabling us to visualize a sigmoidal activation curve for individual astrocytes as a function of stimulation strength. We found that the inflection points of these activation functions varied across cells (Figure 3H), reflecting a lower or higher sensitivity of individual cells. To quantify astrocytic sensitivity, we computed the ratio of the astrocytic response to an intermediate stimulation (R5Hz), normalized by its response to a strong stimulation (R20Hz) that we consider as an approximation of a saturating stimulus: S5/20 = R5Hz/R20Hz (Figure 3I, top). Interestingly, this cell-specific sensitivity was a stable property across days for individual astrocytes (Figure 3I, bottom). These results show that the diversity of astrocytes within hippocampal CA1 spans a large range of response sensitivities for LC stimulation, and that these cell-specific functional properties are stable and consistent within and across days.

### Astrocytic activity during natural arousal diverges from LC-evoked responses

With astrocytes activated by both natural arousal and LC stimulation (Figure 1J-N), one might assume that cells which are more strongly activated by natural arousal and body movement would also be more strongly activated by LC stimulation. However, we did not find evidence for such a relationship (Figure 3J), indicating that the axes of natural arousal and LC stimulation align only weakly. Using sensitivity instead of LC response magnitude, we found a positive albeit relatively weak correlation with movement modulation (Figure S3-1), indicating at least a partial alignment of astrocytic movement modulation with the sensitivity to LC stimulation.

To further investigate whether natural arousal and LC stimulation evoke aligned or distinct astrocytic responses, we compared the similarity of astrocytic population patterns across different contexts: episodes of natural arousal with increased body movement, or LC stimulation of varying intensities (5 Hz or ≥20 Hz). First, to characterize population activity patterns, we sorted recordings according to the cellular response strength to LC stimulation. Interestingly, this sorting did not reveal any obvious alignment of responses to natural arousal during movement (Figure 3K). To quantify this observation, we systematically identified global astrocytic activation patterns either during elevated body movement or upon LC stimulation (see Methods) and computed the similarity between patterns during the same (e.g., movement vs. movement) or different conditions (e.g., movement vs. LC stimulation). We found a striking similarity of population patterns among different instances of movement, in agreement with recent findings in cortex (Fedotova et al., 2025), and among different instances of LC stimulation. However, we observed only low similarities between the two groups (Figure 3L). These analyses based on population activity patterns reinforce the idea that there is only weak alignment between astrocytic responses to arousal and to LC stimulation on a cellular basis, despite the similarity of the bulk response (Figure 1J-N).

### Pyramidal cells exhibit non-specific, uniform inhibition in response to LC stimulation

We found that pyramidal cells were inhibited upon LC stimulation (Figure 1Q-S; Figure 2K), but this effect emerged only after averaging across neurons. We therefore wondered whether individual pyramidal cells might display heterogeneous response profiles, potentially reflecting differences in cell identity or pre-stimulation activity state. At the single-cell level, pyramidal cell activity initially appeared largely unsorted (Figure 2I), but visible patterns became apparent when cells were reordered by their LC-evoked activity change (Figure 4A) or by the timing of their activity peaks (Figure 4B). The stimulus-aligned representation (Figure 4A) suggests that LC stimulation results in both inhibited neurons, consistent with fiber photometry results (Figure 1Q-S), as well as a set of activated neurons. However, it is known that pyramidal cells in hippocampal CA1 are sequentially active even in the absence of external stimulation, a phenomenon described as “time cells” (Eichenbaum, 2014). Sorting by sequential time bins indeed revealed this intrinsic feature of pyramidal cell activity (Figure 4B), and also highlights that fewer neurons are preferentially active in the sequential time bins following LC stimulation (Figure 4C,D). These results indicate that the observed apparent activation of pyramidal cells is a residual signature of ongoing sequential dynamics, rather than a direct effect of LC stimulation.

**Figure 4.**
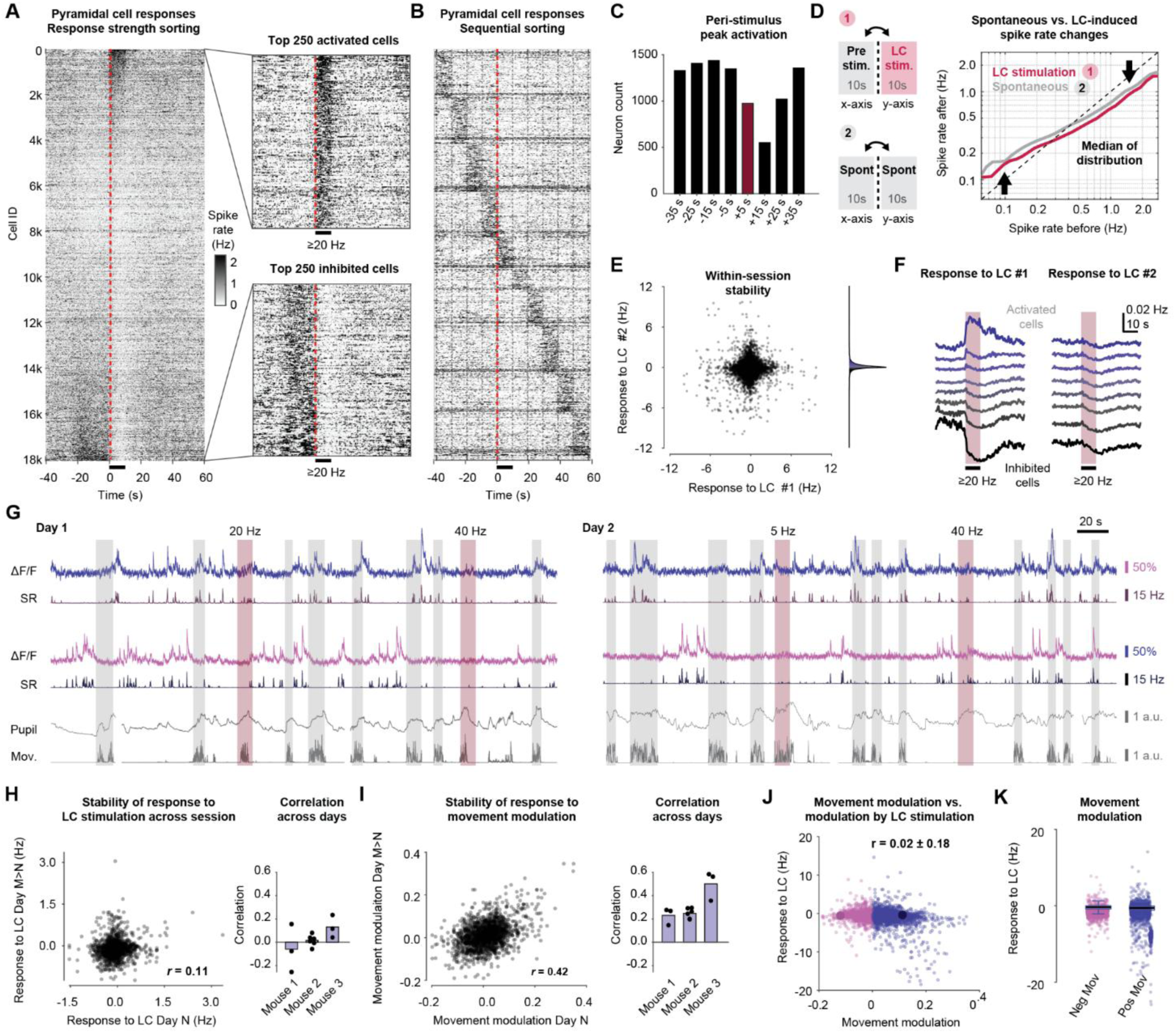
Pyramidal cells exhibit distinct and stable response patterns during natural arousal but respond uniformly to LC stimulation. **A**, All cellular responses to ≥20 Hz LC stimulation, shown as the spike rate estimated from ΔF/F traces and sorted by the spike rate change during the 10 s stimulation window compared to the 20 s before stimulation, with subselection of the 250 most and least responsive cells. Data pooled from 15 sessions across 5 mice. **B**, Same as in (A) but sorted by the highest activity in 10-s time bins for 60 s before and 90 s after LC stimulation. Neurons with maximum activity outside the displayed time window are not shown. The visualization illustrates the sequential activity patterns of pyramidal cells. **C**, Quantification of number of neurons in each time bin in panel (B). The number of neurons activated in the 10-s bins following LC stimulation is lower than for any other time bins. **D**, Change of spike rate from a 10-s pre-stimulus window (x-axis) to the 10-s window of LC stimulation (y-axis). Spike rate changes are shown both for random time points (spontaneous, gray) and 20-Hz LC stimulation time points (red). Compared to spontaneous spike rate changes, LC stimulation induces a reduction of spike rates. **E**, *Left:* Response to two strong LC stimulations within a single session (stimulation 1: x-axis, stimulation 2: y-axis) to investigate within-session stability. *Right:* Marginal distribution of LC stimulation responses, color-coded as in panel (C). **F**, Analysis of stability of pyramidal cell responses within sessions. *Left:* Cellular responses to stimulation #1, sorted by most to least responsive cells and plotted as the quantiles shown in panel (C, right). *Right:* The responses of the same cells to stimulation #2 within the same session, with the sorting maintained from stimulation #1. No evidence for response stability can be observed. **G**, Example recording of negatively-movement modulated (top) and positively-movement modulated pyramidal cell (below), together with behavioral proxies for arousal (pupil diameter, body movement; bottom). Raw ΔF/F traces (light color) as well as estimated spike rates (dark color) are shown, matching the color-code in panels (J,K). Episodes of natural arousal encompassed by body movement are highlighted with gray shading (manual annotation). Episodes of optogenetic stimulation of LC are highlighted with red shading.The same pyramidal cells are tracked for day 1 (left) and day 2 (right) **H**, *Left:* Analysis of stability of pyramidal cells responses towards LC stimulation across sessions (day N vs. day N+1), pooled across all sessions and animals. *Right:* Correlation of responses across days for each animal and each session pair. **I**, *Left:* Analysis of stability of pyramidal cell activity locked to body movement across sessions (day N vs. day N+1), pooled across all sessions and animals. *Right:* Correlation of responses across days for each animal and session pair. **J**, Comparison of the cellular responses to natural arousal (body movement) vs artificial arousal (LC stimulation), not exhibiting any visible correlation. Cells modulated positively vs. negatively by body movement are color-coded. The two neurons shown in (G) are highlighted with large dark symbols. **K**, LC response magnitude of positively-modulated (−0.5 ± 1.9 Hz; median response ± S.D. across neurons). vs. negatively-movement modulated cells (−0.3 ± 1.1 Hz). Color-code as in (J).

Next, we examined how LC stimulation influences the ongoing sequential dynamics of hippocampal pyramidal cells. One possibility is that LC activity amplifies existing neuronal patterns by increasing the activity of already-active neurons while further suppressing less-active ones, consistent with ideas of an LC-mediated increase of neuronal signal-to-noise (Berridge and Waterhouse, 2003; Bouret and Sara, 2002). Using our single-cell two-photon imaging dataset, we quantified the change of neuronal activity upon LC stimulation as a function of pre-stimulus activity (Figure 4D). Interestingly, we find that the net inhibitory effect of strong LC stimulation (≥20 Hz) was uniform across all levels of pre-stimulus activity (Figure 4D; Figure S4-1A,B). The same pattern was observed with moderate stimulation (5 Hz) (Figure S4-1C). These analyses indicate that LC stimulation does not selectively modulate neurons according to their current activity state. Instead, it appears to exert a relatively uniform influence on pyramidal cells, irrespective of their ongoing activity.

Next, we asked whether cell identity determines the strength or direction of a neuron’s response to LC stimulation. We found that neurons that were inhibited (or seemingly activated) by LC stimulation did not show consistent response characteristics across repeated LC stimulations within the same imaging session (Figure 4E,F). Consistent with this observation, patterns of activation or inactivation by LC stimulation were not stable across days when neurons were tracked across sessions (Figure 4G,H). These results further support the idea that pyramidal cells do not exhibit consistent cell-specific responses to LC stimulation. Instead, activation of LC appears to produce a broad, non-specific inhibition that is distributed across the population without targeting particular pyramidal cell subpopulations.

To demonstrate that activity patterns of pyramidal cells were indeed meaningful despite the absence of specific responses to LC, we confirmed that the positive or negative modulation by body movement observed earlier (Figure S2-4) was maintained also across days (Figure 4I). This stability suggests that specific pyramidal cell populations consistently respond to natural arousal (see also the example neurons in Figure 4G). However, neurons that were positively modulated by movement were not more likely to be positively modulated by LC stimulation (Figure 4J,K; response to LC stimulation, median ± S.D.: −0.5 ± 1.9 Hz for positively modulated and a significantly higher −0.3 ± 1.1 Hz for negatively modulated neurons; p < 10^−10^, Wilcoxon rank sum test). This analysis indicates that the cell-specific activity patterns of pyramidal cells in the hippocampus during arousal and movement are not mediated by LC activity.

### Interneurons exhibit stable responses to LC stimulation and arousal

For interneurons, the cell-averaged responses exhibited two main temporal response components to LC stimulation that were particularly apparent after high-intensity stimulation (≥20 Hz): a transient activation, followed by a prolonged inactivation (Figure 2O). To characterize these responses at the single-cell level, we visualized interneurons activity evoked by strong stimulation (≥20 Hz), pooled across all sessions. When we sorted responses by the change in neuronal activity during the 10-s stimulation window (Figure 5A, 3970 responses from 1168 neurons, 10 sessions, 3 mice), we observed a continuum of response types, ranging from strongly activated neurons (Figure 5A, right/top) to spontaneously active interneurons that were inhibited by LC stimulation (Figure 5A, right/bottom). Similar effects for these and other analyses were observed in an additional animal which was analyzed separately due to prominent movement evoked by LC stimulation (Figure S5-1; 2378 responses, 561 neurons, 4 sessions); all further analyses, unless otherwise stated, were performed on animals without this movement confound.

**Figure 5.**
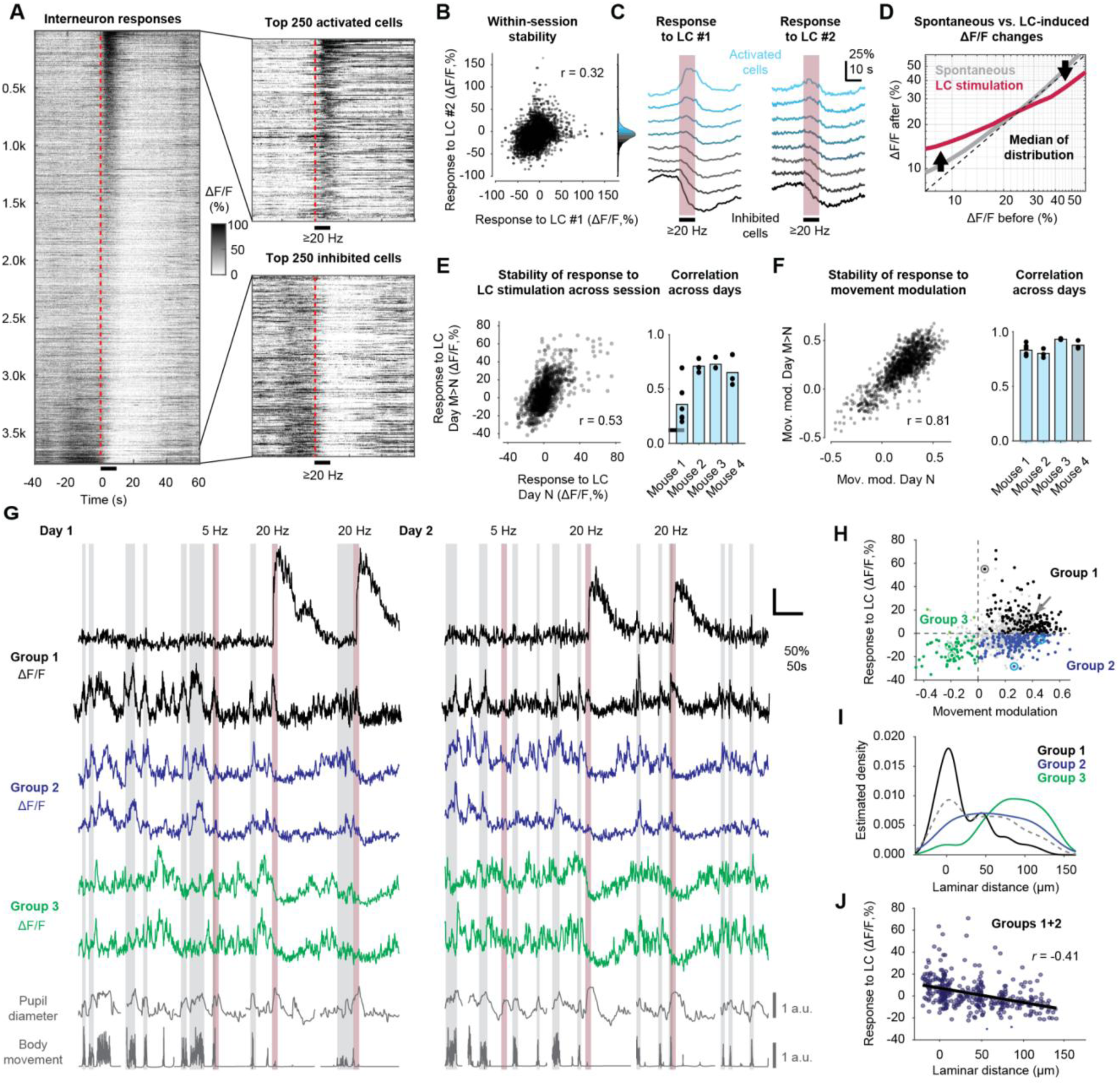
Distinct interneuron populations across laminar position within deep CA1 in response to movement and LC stimulation. **A**, All interneuron responses to ≥20 Hz LC stimulation, sorted by the response during the 10-s stimulation window, with subselection of the 250 most and least responsive cells. Data pooled from 9 sessions across 3 mice. One mouse was excluded due to prominent LC stimulation-related movement (shown separately in Supplementary Figure S5-1). **B**, *Left:* Response to two strong LC stimulations within a single session (stimulation 1: x-axis, stimulation 2: y-axis) to investigate within-session stability. *Right:* Marginal distribution of LC stimulation responses, colorcoded as in panel (C). **C**, Analysis of stability of interneuron responses within sessions. *Left:* Cellular responses to stimulation #1, sorted by most to least responsive cells and plotted as the quantiles shown in panel (C, right). *Right:* The responses of the same cells to stimulation #2 within the same session, with the sorting maintained from stimulation #1. **D**, Change of spike rate from a 10-s pre-stimulus window (x-axis) to the 10-s window of LC stimulation (y-axis) as in Figure 4D. Spike rate changes are shown both for random time points (spontaneous, gray) and 20-Hz LC stimulation time points (red). Compared to spontaneous spike rate changes, LC stimulation tends to activate silent and suppresses active interneurons. **E**, *Left:* Analysis of stability of interneuron responses towards LC stimulation across sessions (day N vs. day N+1), pooled across all sessions and animals. *Right:* Correlation of responses across days for each animal and each session pair. The right-most bar plot, here and in the next panel, corresponds to the animal described in Supplementary Figure S5-1). **F**, *Left:* Analysis of stability of interneuron activity locked to body movement across sessions (day N vs. day N+1), pooled across all sessions and animals. *Right:* Correlation of responses across days for each animal and session pair. **G**, Examples of distinct interneuron response types, 2 exemplary neurons for each group described in panel (H). Episodes of natural arousal encompassed by body movement are highlighted with gray shading (manual annotation). Episodes of optogenetic stimulation of LC are highlighted with red shading. Simultaneously recorded body movement and pupil diameter as proxies for natural arousal are shown below. The same interneurons are shown for day 1 (left) and day 2 (right). **H**, Relationship of natural arousal (x-axis, quantified by body movement modulation of each cell) and response to LC stimulation (y-axis). Upon discarding non-reliably responsive cells (gray), three primary groups remain: Group 1, which responds positively to both LC stimulation and movement (black); Group 2, which responds positively to movement but is primarily inhibited by LC stimulation (blue); and Group 3, which is inhibited both by LC stimulation and negatively modulated by body movement (green). **I**, Laminar distribution within deep CA1 of the three cell groups shown in panel (H). “Zero” indicates the position of the pyramidal cell layer, with positive distance going deeper into the *stratum oriens*. Neurons responsive to LC stimulation (black, Group 1) are located closer to the pyramidal cell layer. Neurons negatively modulated by movement and LC stimulation (green, Group 3) are primarily located in deep layers close to the corpus callosum. Neurons from Group 2 (blue) are distributed across all recorded laminae. The overall distribution of recorded interneurons is shown in the background as a gray dashed line. **J**, Neurons responding positively to movement (Groups 1 and 2) increase their response strength to LC stimulation when located more superficially, i.e., closer to the pyramidal cell layer.

To assess the stability of interneuron response types, we systematically compared responses across LC stimulations. We observed that interneurons responded consistently across repeated stimulations within the same session (≥20-Hz stimulations; Pearson’s correlation coefficient, *r* = 0.32, p < 10^−10^; Figure 5B). In line with that, the sorting of neurons from activated to inhibited was preserved across stimulations within a session (Figure 5C). Although responses to moderate LC stimulations (5 Hz) were weaker than those to strong LC stimulations (≥20 Hz) (Figure 2N), the response profiles across neurons were similar (correlation r = 0.31; Figure S5-2), suggesting that effects seen for strong LC stimulations are likely to generalize also to regimes of weaker LC activation, for which our experimental data are noisier due to smaller response amplitudes.

Next, we investigated whether single-cell responses to LC stimulation could be predicted by pre-stimulation activity. We observed that interneurons with lower pre-stimulation activity tended to be activated by LC stimulation, whereas those with higher activity were typically inhibited (Figure 5D; Figure S5-3). This effect disappeared when random episodes of spontaneous activity without stimulation were analyzed (Figure 5D). This dependence of LC stimulation responses on pre-stimulation levels for interneurons could be driven by either the transient state of a cell prior to stimulation, or by inherent cell identity. To distinguish between these possibilities, we permuted trial identity for pre-stimulation levels. The effect persisted despite the permutation control (Figure S5-3B). These analyses indicate that, while response magnitude can be explained by pre-stimulus activity, it is equally well and more parsimoniously explained by cellular identity.

Tracking cells across sessions, we found that LC-evoked response patterns of interneurons remained relatively stable across days for all animals (Figure 5E, correlation *r* = 0.53). In addition, we quantified the modulation of individual cell responses by body movement and observed an even higher across-session stability (Figure 5F, *r* = 0.81). Together, these results demonstrate that individual interneurons respond in a consistent and cell-specific manner to both natural arousal and optogenetic LC stimulation.

### Dissociation between natural arousal and LC modulation in interneurons

We next asked whether interneurons modulated in a specific manner by body movement exhibited similar responses to optogenetic activation of the LC. If the LC were a significant driver of arousal-related modulation of hippocampal interneurons, movement-related and LC-evoked responses should align for at least a subset of interneurons. However, such an alignment was not visible in interneurons (Figure 5G).

To analyze this relationship more systematically, we first considered the potential confounding effect of modulation by concomitant movement during episodes of LC stimulation. We modeled each interneuron’s activity as a function of past body movement using dilated linear regression and subtracted the neuronal activity predicted from movement alone (Methods; Figure S5-4). All subsequent analyses of responses to LC stimulation use the resulting activity traces where the effect of movement is regressed out.

When comparing LC stimulation responses and movement modulation, we noticed that many interneurons responded to movement or LC stimulation, but with only weak correlation between the two metrics at the level of individual interneurons, and the remaining variability seemed to be dominated by subgroup effects rather than a systematic correlation (Figure 5H, correlation *r* = 0.26 across 1,149 neurons). Since residual movement-related responses sometimes overshadowed small responses to LC stimulation, we restricted our further analysis on cells with responses that were reliable across stimulations (see Methods; colored dots in Figure 5H). These reliably responding cells were found across all animals and recording sessions (Figure S5-5). These reliably responding interneurons fell into three groups: Group 1 and 2 were both positively modulated by body movement, with Group 1 (black in Figure 5H) being activated by LC stimulation and Group 2 (blue in Figure 5H) inhibited. Group 3 interneurons (green in Figure 5H) were negatively modulated by body movement and therefore active during rest; these interneurons were also strictly and consistently inhibited by LC stimulation. Interestingly, interneurons within Group 1 that exhibited the strongest activation by LC stimulation were only weakly or not at all modulated by movement (Figure 5G,H). Together, these results show that both body motion (as a proxy for natural arousal) and optogenetic LC activation drive consistent response patterns in hippocampal interneurons, but the two forms of modulation often evoke different responses within the same cells, indicating only weak alignment between LC-driven and arousal-driven interneuron activity.

### Laminar organization of LC-responsive interneurons

Our interneuron imaging experiments spanned the deep portion of CA1, encompassing *stratum oriens*, the *stratum pyramidale*, and the deep part of *stratum radiatum*. Within this part of CA1, we found that the functionally defined interneuron groups showed a non-random laminar distribution. Interneuron activity patterns were more similar *within* laminae and more distinct *across* the layers of deep CA1 (Figure S2-4), motivating us to investigate the logic of how functionally defined interneuron types (Figure 5H) are distributed along the laminar axis of deep CA1.

To enable the mapping onto this axis, we acquired *in vivo* 3D two-photon imaging stacks encompassing the imaged planes, and measured each interneuron’s laminar distance from the center of the pyramidal cell layer (Methods; Figure 5I). We found that interneurons activated by LC stimulation (Group 1) were strongly enriched in the superficial part of *stratum oriens*, close to the pyramidal cell layer (median laminar distance [IQR]: 6.4 μm [0.0‒45.5 μm]). In contrast, interneurons inhibited by LC stimulation and negatively modulated by body movement (Group 3) tended to be located in deeper layers of *stratum oriens,* closer to the corpus callosum (median [IQR]: 89.8 μm [65.4‒112.6 μm]). Finally, group 2 interneurons showed no clear laminar preference (median [IQR]: 52.3 μm [22.2‒95.0 μm]).

The distinction between Group 1 and Group 2 interneurons may also be described by a continuum rather than discrete response types to LC stimulation. We therefore quantified the response magnitude to LC stimulation as a function of laminar distance across neurons from groups 1 and 2 (Figure 5J; Figure S5-1C for the outlier mouse). This analysis revealed a continuous decrease of activation by LC stimulation as a function of laminar distance from the pyramidal cell layer (*r* = −0.41, p < 10^−10^, n = 383 cells). In contrast, movement modulation exhibited only weak, if any, laminar dependence for the same set of cells (*r* = −0.13, p = 0.01; Figure S5-6). This finding provides further evidence for a dissociation between LC-evoked and arousal-related interneuron activity, based on the distinct laminar location of their somata.

Together, our results show that hippocampal interneurons in deep CA1 exhibit stable and consistent responses to both natural arousal and optogenetic LC stimulation. Additionally, we further find that functionally distinct LC-response types are spatially segregated by a logic that follows the laminar organization of deep CA1.

### Activation of cell types across stimulation intensities

Throughout our experiments we noticed that astrocytes appeared to be particularly responsive even to weak LC stimulation. To enable a direct comparison, we visualized average responses as a function of LC stimulation intensity (Figure 6). Previous analyses showed that a subpopulation of interneurons responded early during the 10-s stimulation (Figure 5), whereas broad inhibition of neurons occurred primarily in the 10-s window after stimulation (Figure 4 and 5). We therefore analyzed the response magnitudes and population activity patterns separately for these two time windows (‘early’ and ‘late’, Figure 6B,E,H vs. Figure 6C,F,I).

**Figure 6.**
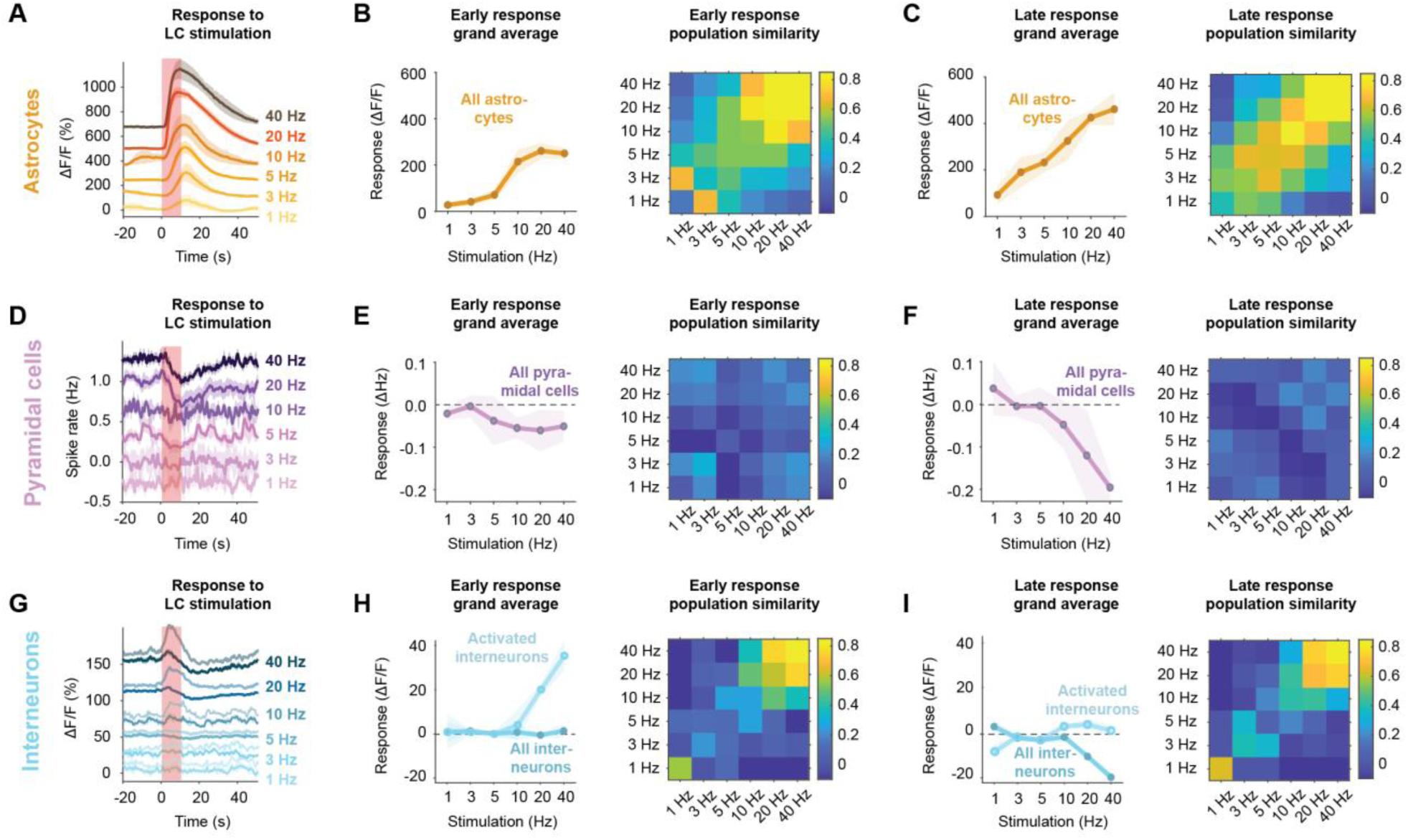
– Cell-type-specific response profiles to LC stimulation across intensities. A, Mean astrocytic population responses to optogenetic LC stimulation at increasing frequencies (1-40 Hz) for astrocytes. Shaded region indicates stimulation period. Responses are offset on the y-axis to improve readability. **B**, Astrocytic response properties during the early response window (10-s window during stimulation) for astrocytes. *Left:* grand-average response magnitude. *Right:* population similarity across stimulation intensities computed with Pearson’s correlation of the population activity vector. **C,** Same as in (B) but for the late time window (10 s after stimulation). **D-F,** Same as in (A-C) but for pyramidal neurons. **G-I,** Same as in (A-C) but for interneurons. In (G-I), the subpopulation of interneurons that were activated by stimulation intensities ≥20 Hz are shown separately (lighter color) together with the entire population (darker color). The similarity measurements are based on the entire interneuron population. For all panels, values are averaged for astrocytes across 495 cells, with 6 stimulations from 3 sessions and 3 animals for 1 Hz, 495/6/3/3 for 3 Hz, 2207/34/12/4 for 5 Hz, 495/7/3/3 for 10 Hz, 2207/32/12/4 for 20 Hz, and 495/6/3/3 for 40 Hz. For interneurons, 368/7/3/3 for 1 Hz, 368/6/3/3 for 3 Hz, 1710/41/14/4 for 5 Hz, 368/6/3/3 for 10 Hz, 1710/39/14/4 for 20 Hz, and 751/12/6/3 for 40 Hz. For pyramidal cells, 672/4/2/2 for 1 Hz, 672/4/2/2 for 3 Hz, 4425/37/14/6 for 5 Hz, 672/4/2/2 for 10 Hz, 4425/36/14/6 for 20 Hz, and 3457/17/9/3 for 40 Hz.

We found that astrocytes exhibited clear responses even at low stimulation intensities (Figure 6A). During the early 10 s-response window, while stimulation was still ongoing, stronger stimulation was necessary to trigger a response (Figure 6B), likely because of the slow and delayed activation of somata (Rupprecht et al., 2024). In the 10 s-time window after stimulation, however, the population response of astrocytes increased monotonically with stimulation intensity (Figure 6C). At the population level, response patterns were similar for comparable stimulation intensities but diverged across distinct intensities (Figure 6C, right). Such an effect cannot arise from a uniform increase in responses across individual astrocytes, and indicates that additional cells were recruited at higher intensities, while others saturated, consistent with earlier analyses (Figure 3H–I). This distinction would not be apparent from population-level recordings such as fiber photometry.

For neurons, the inhibitory effect of LC stimulation emerged only at high stimulation intensities (≥ 10 Hz) and primarily during the late response window (Figure 6D-I). Stable population response patterns were observed for interneurons but not for pyramidal cells (Figure 6H,I vs. Figure 6E,F), consistent with single-cell analyses (Figure 4E,F,H and Figure 5C,E). We further analyzed interneurons that exhibited a positive response to strong LC stimulation (≥ 10 Hz) (brighter lines in Figure 6G-I). This subpopulation was activated for stimulation intensities down to 10 Hz, but not for 5 Hz or below.

Together, these analyses suggest that LC activity at low and moderate stimulation intensities primarily engages astrocytes, while effects on neuronal activity – both selective activation and unspecific inhibition – emerge only at higher levels of noradrenaline release in hippocampal CA1.

## Discussion

In this study, we establish the response profiles of the main cell types in hippocampal CA1 to LC stimulation and how these differ from responses to natural arousal. Using optogenetic LC stimulation in combination with fiber photometry and two-photon imaging, we observe selectively induced NA release at levels comparable to arousal and stress, while avoiding direct engagement of other arousal-associated circuits. Our results show that astrocytes are highly sensitive mediators of LC activity in the hippocampus, whereas pyramidal cells and interneurons respond only at higher levels of LC activity. In addition, our results reveal a striking dissociation at the cellular level: all examined cell classes - pyramidal cells, interneurons, and astrocytes - exhibit markedly different response profiles to natural arousal and LC stimulation (Table 1; Figure S6-1). Finally, we identify functionally distinct interneuron subpopulations that exhibit diverse yet consistent responses to LC stimulation and arousal, and are distributed along the laminar axis of deep CA1. Together, our findings reveal the cell-type-specific organization of NA-induced signaling in CA1 and highlight a predominant role for astrocytes in mediating LC-driven modulation in awake mice.

**Table 1.** Overview of functionally defined response characteristics of the investigated cell types. Subgroups are defined based on different response polarities to arousal or LC stimulation (third and fourth column). Accordingly, there are no subgroups for astrocytes, although astrocytes exhibit a spectrum of sensitivities to LC stimulation. Laminar location is defined as “deep” (more dorsal in dorsal CA1) and “superficial” (more ventral in dorsal CA1). “Aligned with population average” indicates whether the overall response directionality (activation vs. inhibition) was the same between responses to arousal and responses to LC stimulation. “Aligned on cellular level” indicates whether the single-cell response profiles were aligned across arousal vs. LC stimulation. “Response stability” indicates whether single-cell response patterns were stable within and across days.

| Hippocampal cell type | Functionally defined subgroups | Response to arousal | Response to LC stimulation | Laminar location | Aligned with population average | Aligned on cellular level | Response stability |
| --- | --- | --- | --- | --- | --- | --- | --- |
| Pyramidal cells | Group 1 | Activation | Inactivation | N/A | No | No | No |
|  | Group 2 | Inactivation | Inactivation | N/A | Yes | No | No |
| Interneurons | Group 1 | Activation | Activation | Superficial | Yes | Weak | Yes |
|  | Group 2 | Activation | Inactivation | Unspecific | No | No | Yes |
|  | Group 3 | Inactivation | Inactivation | Deep | Yes | Partial | Yes |
| Astrocytes | N/A | Prominent activation | Prominent activation | Unspecific | Yes | Partial | Yes |

### Astrocytes as diverse and highly sensitive responders to noradrenaline

In our *in vivo* experiments, astrocytes consistently responded with robust calcium elevations to both natural stressors and optogenetic LC stimulation, closely matching NA dynamics (Figures 1). This aligns with the well-established role of astrocytes as slow, synchronized trackers of arousal (Dombeck et al., 2007; Nimmerjahn et al., 2009; Rupprecht et al., 2024). Importantly, the calcium elevations evoked in astrocytes were evident even at moderate LC stimulation intensities, at which pyramidal cells and interneurons showed no detectable direct responses (Figures 6). This high responsiveness most likely reflects the high sensitivity of astrocytes to NA via ɑ1-adrenergic receptors (Wahis and Holt, 2021).

Notably, sensitivity to LC stimulation varied within the astrocytic population (Figure 3). At the same time, individual astrocytes exhibited stable response amplitudes and sensitivity, suggesting a functional diversity within astrocytic populations. Whether this diversity reflects a continuum of NA sensitivities, possibly driven by differences in α1-receptor expression, the vicinity of noradrenergic fibers to different astrocytes, or distinct astrocyte subtypes, remains unclear. Future work will be needed to relate these functional differences to recently described anatomical and transcriptional subtypes of astrocytes (Endo et al., 2022; Hennes et al., 2025; Viana et al., 2023).

Together, the high NA sensitivity of astrocytes compared to neurons and their diverse response profile makes astrocytes an ideal candidate to act as the primary mediator of the effects of LC in the hippocampal circuit during arousal and moderate levels of stress.

### Unspecific and broad inhibition of pyramidal cells by LC stimulation

Theories about the effect of NA release propose that NA increases the signal-to-noise ratio by enhancing the activity of specific neuronal ensembles while suppressing others (Berridge and Waterhouse, 2003; Bouret and Sara, 2002). These ideas were inspired by brain slice work demonstrating β-adrenergic increase of excitability of hippocampal pyramidal cells by bath-applied NA (Church et al., 2019; Madison and Nicoll, 1982; Sah and Isaacson, 1995) or by optogenetically evoked endogenous NA release in *ex vivo* preparations (Bacon et al., 2020). However, early *in vivo* studies using electrical LC stimulation reported overall neuronal suppression (Segal and Bloom, 1976). In slices, inhibition of pyramidal cells was shown to require high exogenous NA levels, likely involving activation of presynaptic ɑ2-adrenergic receptors (Bacon et al., 2020), or indirect effects of NA on presynaptic activity mediated by astrocytes (Lefton et al., 2025). In our *in vivo* experiments, LC stimulation produced a broad, slow, and non-specific inhibition in pyramidal cells. The reasons for the differences between *in vivo* and *ex vivo* experiments remain unclear and need to be addressed in future work.

In addition, our dataset enabled us to analyze how the ongoing activity of individual pyramidal cells was affected by LC stimulation. This analysis revealed a lack of cell-specific activation or inactivation patterns (Figure 4E,F,H). In addition, responses to LC stimulation did not depend on the activity level of pyramidal cells prior to stimulation (Figure 4D). It is not clear how these findings can be reconciled with the idea that NA release increases activity only for specific neuronal ensembles (Berridge and Waterhouse, 2003; Mather et al., 2016). Together, these findings challenge the notion that LC-evoked NA release directly increases the excitability of hippocampal pyramidal cells *in vivo*. However, future work should investigate whether NA can have excitatory effects on pyramidal cells under specifically designed behavioral conditions.

### Interneurons as laminarly heterogeneous but reliable responders to LC

Interneurons responded to LC stimulation with substantial heterogeneity across the population, yet individual neurons showed stable response profiles across days. Most interneurons displayed a slow inhibition resembling that of pyramidal cells, but a subset of interneurons exhibited, in addition, a transient activation (Figure 5A). Based on their responses to LC stimulation and their modulation by body movement, we identified three functional interneuron groups. Group 1 was activated by LC stimulation and positively modulated by movement.

Group 2 was inhibited by LC stimulation but still positively modulated by movement. Group 3 was inhibited by LC stimulation and negatively modulated by movement (active during rest).

Group 3 interneurons match prior descriptions of deep-laminar neurons with negative movement modulation (Geiller et al., 2020). While this modulation has been loosely explained by laminar position, an association with specific molecular subtypes remains unclear, as small fractions across all major interneuron subtypes (SST, PV, and VIP) were found to be negatively modulated by movement (Arriaga and Han, 2017; Geiller et al., 2020; Turi et al., 2019). Notably, LC-induced inhibition in our dataset was an even stronger predictor of deep laminar position than negative movement modulation alone (Figure 5I,J). These results suggest that laminar location may be a key determinant of both arousal modulation and NA responsiveness that is complementary to, and in some cases as informative as, the molecular identity of interneurons in hippocampus.

Group 1 interneurons, located near the pyramidal cell layer, are particularly intriguing due to their reliable activation by both LC stimulation and movement. However, while these interneurons seem to respond directly to LC-evoked NA, the receptor mediating this effect remains unclear. Slice studies have shown interneuron activation by NA via ɑ1A-receptors (Bergles et al., 1996; Hillman et al., 2009; Madison and Nicoll, 1988). In our study, we found that astrocytes responded robustly to NA through α1-receptors even at low stimulation intensities (Figures 1 and 3), whereas interneuron activation *in vivo* emerged only at substantially higher intensity (Figures 1, 5, S5-3C). Further complicating the comparison, the activation of inhibitory cells in slices was shown to occur in SST-expressing cells (Hillman et al., 2009, 2005), which, based on newer transcriptomic profiling, express relatively low levels of ɑ1-receptors (Yao et al., 2021; Smith and Von Zastrow, 2022). This mismatch raises the question whether the mechanism underlying interneuron activation observed in slices is related at all to the activation that we observed *in vivo*. Higher expression levels of α1-receptors in CA1 were found in a subset of Lamp5 interneurons and, to a lesser extent, in VIP interneurons (Smith and Von Zastrow, 2022; Yao et al., 2021). Lamp5 interneurons comprise Ivy cells and neurogliaform cells. While neurogliaform cells are located in *stratum moleculare* and hence out of reach for our *in vivo* imaging experiments, Ivy cell somata are located near the pyramidal cell layer (Tzilivaki et al., 2023), matching the laminar distribution of Group 1 interneurons (Figure 5I). VIP subtypes also remain interesting candidates, given their presumed roles in pyramidal cell plasticity during neuromodulation (Turi et al., 2019; Neubrandt et al., 2025) and the role of hippocampal NA in responding to novelty (Heer and Sheffield, 2023; Kaufman et al., 2020; McKenzie et al., 2025). However, the functional heterogeneity within VIP populations (Dellal et al., 2025) and the challenges of selectively targeting specific interneuron subtypes *in vivo* render a definitive identification difficult.

These inconsistencies between slice physiology, receptor expression profiles, and *in vivo* responsiveness highlight the need for further experiments combining functional imaging and more precise transcriptomic classification to accurately map NA-responsive interneurons to molecularly defined subtypes. At present, our functional grouping and its association with laminar organization in deep CA1 provides a starting point for understanding how hippocampal interneurons respond to endogenous NA release.

### Mechanisms and physiological relevance of broad neuronal inhibition

The broad inhibition of both pyramidal cells and most interneurons in response to LC stimulation is an observation that is both striking and puzzling. Given that interneurons regulate pyramidal cell excitability, one might expect that silencing interneurons would lead to disinhibition of pyramidal cells; instead, we find that both populations are suppressed. Several explanations could account for this observation. One possibility is that the LC acts indirectly through other brain regions that produce an effective silencing of CA1 independent of local NA release. Another is that the subset of interneurons that was transiently activated by LC stimulation (group 1 in Figure 5) is sufficient to induce a sustained suppression of local activity of both pyramidal cells and interneurons. Alternatively, inhibition may be mediated by an alternative local mechanism not dependent on interneurons. Interestingly, the slow dynamics of the inhibitory response seems to mirror the similarly slow time course of NA release or astrocytic activity (Figures 1 and 2). This similarity puts forward the hypothesis that the observed neuronal inhibition could be mediated, at least partially, either directly by the local action of NA, or indirectly via astrocytes. Astrocytes express a diverse range of adrenergic receptors, particularly α1-adrenergic receptors, which are known to promote intracellular calcium signaling. Experiments in *ex vivo* slice preparations and in zebrafish show that astrocytic activation can indeed exert inhibitory effects on neuronal activity, potentially through the release of gliotransmitters. ATP, for example, is released by astrocytes and metabolized into adenosine, which acts as an inhibitor of neuronal activity via A1 receptors (Chen et al., 2025; Lefton et al., 2025). It remains to be seen whether this signaling pathway can explain the inhibition observed in our experiments.

Irrespective of the exact mechanism, the broad inactivation of both pyramidal cells and interneurons raises the question about its physiological relevance. This wide-spread inhibition is reflected on the behavioral level by the finding that high frequency stimulation of LC can induce behavioral arrest (Carter et al., 2010), in agreement with our own observations (Figure S2-3B). In this light, our results may reflect a mechanism by which LC activation under highly stressful conditions selectively silences hippocampal circuits that are primarily important for cognitive processes. This interpretation aligns with recent ideas that NA may help dampen activation of neuronal circuits during heightened arousal (Reitman et al., 2023). Indeed, LC-evoked NA release has been shown to increase activity of other brain regions such as the lateral habenula (Xin et al., 2025) or periaqueductal gray (Petersen et al., 2025), which are closely connected to escape, aversion and defensive behaviors. Interestingly, a similar suppressive effect of another neuromodulator, serotonin, has been recently described in the visual cortex (Azimi et al., 2020), suggesting that widespread inhibition of neuronal activity might be a more general mechanism of action by multiple neuromodulators. Probing these ideas will require stimulations of neuromodulatory areas combined with recordings in other brain regions using approaches capable of disentangling population averages from single-neuron activity.

To understand the physiological meaning of the observed effects of LC stimulation, it is essential to relate LC stimulation intensities to NA levels during natural behavior. We used behavioral stress paradigms to calibrate the optogenetic stimulation of the LC. Based on this calibration, LC stimulation of moderate intensity (5 Hz) corresponded to an already significant acute stressor, whereas stronger stimulation (20–40 Hz) induced NA levels comparable to those triggered by very high stress (Figure 1). In our experiments, intermediate stimulation strength evoked only modest neuronal responses, prohibiting detailed analyses on the single-cell level due to low signal-to-noise. Consequently, we focused our single-cell analyses on responses to strong LC stimulation (Figure 3 to 5). However, we noticed that activity patterns in response to LC stimulation were qualitatively similar for strong and intermediate stimulation strength (Figure S4-1; Figure S5-3), suggesting that the observed effects likely occur at lower levels of NA as well, albeit with reduced magnitudes.

### Disentangling the effects of natural arousal and LC activity

The LC is the principal source of NA in the brain and a central regulator of arousal. While our optogenetic paradigm successfully reproduced natural NA release dynamics, astrocytic and especially neuronal responses differed significantly between the two conditions. To compare these states, we identified arousal episodes using body movement during head-fixation as a well-established behavioral proxy (McGinley et al., 2015; Stringer et al., 2019; Collins et al., 2023). Although it would be interesting to investigate more intense stressful conditions (cf. Figure 1) with single-cell resolution, such behaviors are not as easily compatible with head-fixed behaviors and high-quality two-photon imaging.

In our head-fixed experiments, individual astrocytes were reliably activated under both conditions (Figure 1; Figure 3). However, astrocytes strongly activated during arousal were not likely to be equally sensitive to LC stimulation, resulting in distinct population activity patterns (Figure 3J-L). This difference, although apparently minor, indicates that LC-evoked NA release does only partially replicate the natural effect of NA release during arousal and points out a gap of understanding that future work should fill. For example, it would be important to assess astrocytic responses to different neuromodulators (including NA, acetylcholine, serotonin) and neurotransmitters (GABA, glutamate), and their expression profiles of the respective receptors.

For pyramidal cells and interneurons, the misalignment between the two conditions was even more pronounced. On average, neurons were activated during arousal but inhibited during LC stimulation (Figures 1 and 2). Pyramidal cells exhibited robust positive or negative modulation by body movement, but no such consistent modulation by LC stimulation (Figure 4). Among interneurons, a subset displayed matching response polarity across conditions (Groups 1 and 3 in Figure 5H), but response magnitudes to arousal and LC stimulation were not correlated within these groups. In addition, further supporting the dissociation from an anatomical perspective, responsiveness to LC stimulation was organized along the laminar axis in deep CA1 (Figure 5J), while modulation by arousal was not.

Together, these findings highlight the low similarity between neural activity during LC stimulation and natural arousal. This difference may be partially explained by technical factors such as non-natural activation patterns of LC neurons by optogenetics, but is also very likely to reflect that other neuromodulators or neurotransmitters are released during arousal, resulting in a cellular response profile that is not aligned with LC-evoked NA release alone. Therefore, our results suggest that LC activity and NA release, while central players for arousal, are not the primary driver of the neuronal dynamics in the hippocampus concurrent with arousal, and may either lower background signal to enhance signal-to-noise for relevant incoming stimuli (Berridge and Waterhouse, 2003), or could counterbalance other neuromodulatory inputs.

For instance, acetylcholine (ACh), released from the basal forebrain and medial septal projections, is another key neuromodulator implicated in arousal, attention, and locomotion (Muñoz et al., 2017; Bugeon et al., 2022). ACh release correlates with pupil diameter, a common proxy for arousal (Reimer et al., 2016) and has been demonstrated to drive locomotion-related activity of cortical interneurons (Fu et al., 2014) and hippocampal pyramidal cells (Fuhrmann et al., 2015). Other neuromodulatory systems such as serotonin, histamine, orexin, and CRH are also linked to arousal state and are likely to shape hippocampal neural activity. A systematic assessment of these systems and their influence on all hippocampal cell types, ideally through selective *in vivo* activation of each pathway in isolation and in a combinatorial manner, would help understand the components of the arousal system and their interaction. Our findings underscore that, while LC-driven NA release produces consistent and cell-specific responses, additional neuromodulatory or circuit-level mechanisms are necessary to produce the full phenotype of arousal-related activity in hippocampal astrocytes, pyramidal cells, and interneurons.

## Contributions

S.N.D., J.B. and P.R. conceived the study. S.N.D, A.M., and R.Z. performed the fiber photometry experiments, S.N.D. and P.R. performed two-photon experiments. M.W. established fiber photometry analysis pipeline and S.N.D. and M.W. analyzed fiber photometry data. S.N.D. and P.R. performed and analysed two-photon imaging experiments. S.N.D. and P.R. conceptualized the results and wrote the original draft, S.N.D., A.M. M.W., R.Z., F.H., J.B., and P.R. reviewed and edited the manuscript draft. J.B., F.H., and P.R. supervised the study and acquired the funding.

## Acknowledgements

The group of J.B. was supported by the ETH project grant ETH-20 19-1, SNSF grants 310030_172889 and 310030_204372, the Botnar Research Center for Child Health, the Swiss 3R Competence Center, Roche and the Hochschulmedizin Zürich Flagship project STRESS. The group of F.H. and P.R. was supported by grants from the Swiss National Science Foundation (project grant 310030B_170269 to F.H.; SNSF Advanced Grant TMAG-3_216336 to F.H.; Ambizione Grant PZ00P3_209114 to P.R.). We thank F. Rössler for help with behavioral data analyses. We thank J.-C. Paterna from the VVF of the Neuroscience Center Zürich, a joint competence center of ETH Zürich and the University of Zürich, for producing the viral vectors and viral vector plasmids. We thank S. Ghosh for initial help with calcium imaging preprocessing. We thank the staff of the EPIC for the excellent animal care and their service to our animal facility and J. Bode, J. Leonardi and A. Madhavan for maintaining the animal colony. Figures showing schematics were created with BioRender.com.

## Data and software availability

Preprocessed two-photon calcium imaging and behavioral raw data, metadata and analyzed results are available online at Zenodo (https://doi.org/10.5281/zenodo.18672975). Analysis scripts associated with the manuscript and dataset will be made available before publication in a peer-reviewed journal on GitHub (https://github.com/PTRRupprecht/Duss-et-al-2026).

## Methods

### Animals

All animal procedures were conducted in accordance with the Swiss federal guidelines for the use of animals in research and approved by the Cantonal Veterinary Office of Zurich. Heterozygous C57BL/6-Tg(Dbh-iCre)1Gsc (DBH-iCre) mice (Parlato et al., 2007) were kept in standard housing on a 12 h light/dark cycle.

### Stereotaxic surgery

Virus injection and fiber implantation

Stereotaxic surgery was performed on 2 to 4 months old DBH-iCre mice under 2% isoflurane anaesthesia with a subcutaneous dose of 5 mg/kg Meloxicam and 0.1 mg/kg Buprenorphine, combined with a local anaesthetic (>2 mg/kg Lidocaine and 2 mg/kg Bupivacain). DBH-iCre was chosen since it provides specific and efficient genetic access to LC neurons (Wissing et al., 2025). Animals were placed into a stereotaxic apparatus onto a temperature controlled heating pad (35° C-37° C). Their skull was exposed, bregma was located and the skull placement corrected for tilt and scaling. For virus delivery and optical fiber implantation, a small hole was drilled above the LC and hippocampus. For fiber photometry (n(male) = 9, n(female) = 22) and opto-two-photon (n(male) = 6, n(female) = 7), mice were then injected unilaterally or bilaterally (coordinates: AP −5.4 mm, ML +/-0.9 mm, DV −3.8 mm), with 0.6 - 0.8 µl of an AAV construct carrying the optogenetic construct (for LC stimulation ChrimsonR (ssAAV-5/2-hEF1α/hTLV1-dloxChrimsonR_tdTomato(rev)-dlox-WPRE-bGHp(A)), 4.7×10^12^ vg/ml unilaterally; for LC inhibition bilateral JAWS (ssAAV-5/2-hSyn1-dlox- Jaws_KGC_EGFP_ER2(rev)-dlox-WPRE-bGHp(A)-SV40p(A)), 6.4×10^12^ vg/ml bilaterally; Viral Vector Facility (VVF), Neuroscience Center Zurich) using a pneumatic injector (Narishige, IM-11-2) and calibrated microcapillaries (Sigma-Aldrich, P0549). For hippocampal recordings, mice were injected with 0.2-0.3 µl of a genetically encoded NA sensor (either GRAB_NE1m_, ssAAV-9/2-hSyn1-GRAB(NE1m)-WPRE-hGHp(A), 5.5×10^12^ vg/ml, GRABNE2m, ssAAV-9/2-hSyn1-GRAB_NE2m-WPRE-hGHp(A), 2×10^12^ vg/ml, or nLightG, AAV-9/2- hSyn1-chl-nLightG-WPRE-bGHp(A), 5.5×10^12^ vg/ml; VVF, Neuroscience Center Zurich; coordinates: AP −3.2 mm, ML −3.3 mm, DV −3.8 mm) or 0.4 µl of a genetically encoded calcium sensor (GFAP-GCaMP6s, ssAAV-9/2-hGFAP-hHBbI/E-GCaMP6s-bGHp(A), 5×10^12^ vg/ml; CaMKIIα-GCaMP8m, ssAAV-5/2-mCaMKIIα-jGCaMP8m-WPRE-bGHp(A), 1-2×10^12^ vg/ml; mDlx-HBB-GCaMP8m, ssAAV-1/2-mDlx-HBB-chI-jGCaMP8m-WPRE- SV40p(A), 2×10^12^ vg/ml; VVF, Neuroscience Center Zurich; coordinates: AP −2 mm, ML −1.5 mm, DV −1.4 to −1.25 mm) in the ipsilateral hippocampus. 5 additional GRAB_NE2m_ animals only received hippocampal injection without LC targeting. For control experiments, mice were injected with 0.2-0.3 µl of an AAV virus encoding an enhanced green fluorescent protein (GFP; ssAAV-9/2-hSyn1-EGFP-WPRE-hGHp(A), 5.5×10^12^ vg/ml; VVF, Neuroscience Center Zurich; coordinates: AP −3.2 mm, ML −3.3 mm, DV −3.8 mm).

Subsequently, for fiber photometry animals, an optical fiber was implanted at 200 μm superior to the injection coordinates (for fiber photometry: diameter 200 µm, NA=0.37; Neurophotometrics, USA; for two-photon imaging: low profile, 90°, diameter 200 μm, NA = 0.66; Doric Lenses, Canada). Optical fibers were glued to the skull using a bonding agent (Etch glue, Heraeus Kulzer GmbH) and a UV-curable dental composite (Permaplast, LH Flow; M+W Dental, Germany) and stitches were used as required. The health of the animals was evaluated by post-operative checks over the course of 3 consecutive days and at least twice, animals received additional analgesia (s.c. injection of 5 mg/kg meloxicam and 0.1 mg/kg buprenorphine). Animals for fiber photometry were left to recover for at least two weeks, while animals for two-photon imaging were placed into quarantine for approximately 2 weeks and transferred to another facility where cranial window surgeries were performed. After 3-4 weeks, pupillometry was performed as described below to validate functional expression of the actuator/inhibitor (Privitera et al., 2020). For two-photon experiments, only animals that showed pupil responses upon LC stimulation were subjected to a second surgery for hippocampal window implantation in the same week.

Cranial window

In a second surgery, cranial windows were implanted to enable two-photon imaging of hippocampal CA1. Preparations for this second surgery were identical as described above. In the stereotaxic frame, the sutured skin was reopened for hippocampal window implantation, as described previously (Rupprecht et al., 2024). Briefly, to expose the brain, a 3-mm diameter craniotomy centered at the previous injection site was drilled. For attachment, two layers of light-curing adhesive (Scotchbond Universal, 3M) were applied to the skull and the cement from the first surgery, followed by a ring of dental cement (Charisma, Kulzer) to prevent overgrowth with skin. A 3-mm diameter biopsy punch (BP-30F, KAI) was inserted 1.3 mm into the brain and left in place for >5 min to stop bleeding. With a flatly cut-off injection cannula (Sterican 27G, Braun) connected to a vacuum pump the cortex was carefully removed until the stripes of the corpus callosum became visible. The corpus callosum was left intact. Bleedings were stopped with absorbent swabs (Sugi, Kettenbach) and hemostatic sponges (Spongostan, Ethicon). Then, a cylindrical metal cannula (diameter 3 mm, height 1.2-1.3 mm) attached with dental cement to a 0.17-mm thick coverslip (diameter 3 mm) was inserted into the cavity and positioned with a glass capillary. When no further bleeding occurred, the hippocampal window was fixed in place using dental cement (Tetric EvoFlow, Ivoclar). Tissue glue (Vetbond, 3M) was used to connect the animal’s skin with the ring of dental cement. A head bar was attached using dental cement (Tetric EvoFlow). After surgery, animals were monitored for 3 days with application of antibiotics (2.5% Baytril in drinking water, Vetpharm), and analgesics (Metacam, 5 mg/kg, s.c.; Buprenorphine, 0.1 mg/kg, s.c.)) administered when necessary. Behavioral training started two weeks after surgery. Calcium imaging was performed 2-3 weeks after the start of behavioral training.

### Pupillometry

After surgery, animals were left to recover and for the viruses to express for 3-4 weeks, then pupillometry was performed. Pupil recordings were obtained from the eye ipsilateral to the stimulated LC and simultaneous with anesthetized fiber photometry recordings. Recordings were performed as previously described in (Privitera et al., 2020) with a Raspberry Pi NoIR Camera Module V2 night vision camera, an infrared light source (Pi Supply Bright Pi - Bright White and IR camera light for Raspberry Pi) and a Raspberry Pi 3 Model B (Raspberry Pi Foundation, UK) was used. In short, mice were anesthetized in an induction chamber with isoflurane in a 1:4 O2 to air mixture (4% induction, 2% maintenance). Anaesthesia levels were maintained at 1.5% – 1.75% isoflurane throughout recordings via a breathing mask. Recordings started once pupil diameter had stabilized. Stimulation protocols included 1 min baseline recording followed by 10 s LC stimulation with 2-3 min post-stimulation recording. Inhibition was performed under IR light only and with 30 s laser inhibition.

### Fiber photometry

Recordings during acute stress exposure

Animals were handled for 5 days and patch-cord trained for additional 4-5 days prior to any fiber photometry recordings. All animals first went through a 10 min baseline open field test (data not shown) to get used to the arena (45 cm × 45 cm × 40 cm [L x W x H], black Plexiglas walls and a white PVC floor). The second time in the open field arena, animals were subjected to a mild tail lift stressor. After 3 min baseline recording, animals were lifted by their tail base three times for 10 s and then carefully released. Between each tail lift, animals had 2 min intervals in the open field arena and the experiment ended after 10 min total.

Foot shock experiments were performed in sound insulated and ventilated multi-conditioning chambers (MultiConditioning System, TSE Systems Ltd, Germany, 30 cm x 30 cm x 40 cm [L x W x H]) with dim yellow and IR light. After a 2 min baseline recording, animals received a single 3 s 0.6 mA foot shock from the metal floor grid and post-shock recording continued for another 2 min.

For the cold forced swim stress paradigm, animals were recorded for 2 min in an empty homecage and then placed in a bucket (20 cm diameter, 25 cm deep) of 18 ± 0.1 °C cold water for 3 min minutes. Immediately after stress exposure, mice were returned to their assigned homecage, quickly and carefully dried with a paper towel and left to recover.

Recordings with optogenetic LC stimulation

Optogenetic LC stimulation was performed using a 635 nm laser built into the neurophotometrics fiber photometry system and fiber output power was set to 6 - 10 mW (as measured at the fiber tip when illuminating continuously). For all stimulation protocols, 10-ms pulses of light were applied, but at different repetition frequencies, ranging from 1 Hz up to 40 Hz. Mice were placed in the open field arena for 10 min. After 3 min baseline recording, animals were optogenetically stimulated three times for 10 s with 2 min inter stimulus intervals.

Recordings with optogenetic LC inhibition

Optogenetic inhibition was performed with an external 635 nm laser (10-12 mW power, CNI laser) at 40 Hz for 30 s. For experiments recording NA levels, confirmation of LC inhibition was performed in the open field area. After 2 min baseline recording, the LC was inhibited for 30 Inhibition during the foot shock experiment included 30-s inhibition starting 15 s prior to the shock. and post-inhibition recording continued for 1.5 min. For the tail lift experiment, recording included a 10-s tail lift every 2 min where the 3rd, 4th, 7th, and 8th tail lift were paired with 30-s of inhibition starting 10 s before the tail lift. Inhibition during the foot shock experiment included 30-s inhibition starting 15 s prior to the shock.

Fiber photometry data acquisition

Photometry signals were recorded as previously described in (Grimm et al., 2024) using a commercially available photometry system (Neurophotometrics, Model FP3002) controlled via the open-source software Bonsai (2.6.2 version). The implanted fiber was attached to a pre-bleached recording patch cord (diameter 200 μm, NA = 0.39; Doric Lenses). Two LEDs were used to deliver interleaved excitation light for recording of NA- or Ca²⁺-bound signal (470 nm LED, fluorescent signal: F470) and unbound sensor signal (415 nm LED, fluorescent signal: F415), with 30 Hz sampling frequency for each channel. Excitation power at the fiber tip was set to 25-35 μW.

Quantification and analysis

Analysis of raw photometry data was performed using a custom-written MATLAB script. Change in fluorescence (𝛥𝐹/𝐹) was calculated separately for F^470^ and F^415^:

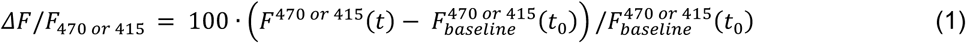

where F^470^ or ^415^(t) signifies the fluorescence value at each time point *t* across the recording and 𝐹^470^ ^𝑜𝑟^ ^415^_*baseline*_(𝑡_0_) denotes the mean value of the signal at baseline (−10 s to 0 s for LC stimulation and foot shock, −10 s to −5 s for tail lift pre-event). To remove movement artifacts, sensor-independent signal (𝛥𝐹/𝐹^415^) were subtracted from sensor-bound signal (𝛥𝐹/𝐹^470^):

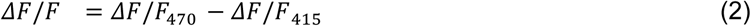

The final signal was smoothed with a 50-point moving mean filter.

### Two-photon microscopy

#### Microscopy setup

A custom-built two-photon microscope was used to monitor calcium signals in astrocytes and neurons in either a single or multiple layers of CA1, similar to previously described (Rupprecht et al., 2024). A femtosecond-pulsed laser (MaiTai, Spectra physics; 905 nm; power below the objective 30-60 mW) was enlarged by a telescope and sent through a scan engine (8-kHz resonant scanner, Cambridge Technology; slow galvo scanner, 6215H, Cambridge Technology) before entering the back aperture of a 10x water immersion objective (Olympus XLPLN10XSVMP, NA 0.6, WD 8 mm), achieving a maximal FOV of 900 µm. Calcium imaging was performed at a rate of approx. 30 Hz (512 x 512 pixels) for a single plane (pyramidal cells) and lower frame rates for multiplane imaging (10 Hz for astrocytes with triple-plane imaging, 7.5 Hz for interneurons with four-plane imaging). Fast z-scanning was performed with a remotely installed electrically tunable lens (Optotune), with axial spacing of adjacent planes by 40 µm. Scanning and data acquisition was controlled with Scanimage 2023 (Pologruto et al., 2003) or, for a small subset of the data, with a custom-written software programmed in C++ (http://rkscope.sourceforge.net/, (Chen et al., 2016)).

#### Behavioral setup for head-fixed two-photon imaging

The behavioral setup was described previously (Rupprecht et al., 2024). The treadmill consisted of two light-weight wheels, one of which was attached to a rotary encoder (Phidgets) to measure locomotion of the animal. A 130 cm long and 5 cm wide velvet belt (88015K1, McMaster-Carr) was equipped with sensory landmarks consisting of self-sticking elements, velcro strips and hot glue. A metal tape attached to a single location of the back side of the belt was used as a reference for an IR sensor to track the animal’s location. IR light (LIU850A, Thorlabs) together with a camera (DMK23UP1300, The Imaging Source, recording at 30 Hz; 16-mm EFL objective MVL16M23, Thorlabs) was used to monitor the animal’s behavior. A UV LED (LED370E, Thorlabs) was directed towards the right eye of the animal, resulting in a preconstriction of the pupil to enable pupil segmentation. Sweetened water rewards (30% sugar) were provided through a metal lick spout at a specific location of the belt. Reward delivery was controlled by a solenoid valve (VDW22JA, SMC). In a subset of experiments, brief air puffs to the left side of the animal’s face were provided randomly and rarely (maximally once per minute).

The behavioral setup was controlled using custom-written Python code, which controlled valves, camera, and microscope acquisition start, and recorded the position of the rotary encoder, the IR sensor and behavioral events like air puffs or water rewards.

#### Optogenetic stimulation

Optogenetic LC stimulation during two-photon calcium imaging was performed using a 635 nm laser (CNI) with a fiber output power of 10 mW (as measured at the fiber tip when illuminating continuously). For all stimulation protocols, 10-ms pulses of light were applied, but at different repetition frequencies, ranging from 1 Hz up to 40 Hz, as for fiber photometry experiments. 5 Hz and 20 Hz were used as standard repetition frequencies for moderate and strong stimulation, respectively. 40 Hz was used as an additional strong stimulation frequency, ensuring that potentially small effects of LC stimulation were maximized for interneuron and pyramidal cell recordings. Analyses for combined 20 Hz and 40 Hz stimulation frequencies are indicated as ≥20 Hz.

#### Behavior training and two-photon imaging experiments

One week before experiments, drinking water of mice was supplemented with citric acid (2% of volume) to motivate the mice during the running task as described before (Rupprecht et al., 2024; Urai et al., 2021). Mice were handled for 15-20 min per day for 3 days and accustomed to the behavioral setup. For the next 3-7 workdays, mice were trained to head-fixation on the treadmill. When animals readily ran on the treadmill and consumed sugar water rewards for 15-20 min, imaging experiments were performed.

A session lasted for 23.6 ± 6.6 min (mean ± s.d.) for awake and 17.6 ± 4.5 min for anesthetized sessions and consisted of 145 s-long segments, spaced by breaks of 5-20 seconds. Sessions recorded on different days maintained the same FOV for most animals (11 out of 13 mice).

### Calcium imaging post-processing

First, rigid movement correction followed by non-rigid movement correction in the xy-plane was applied to raw calcium imaging movies (Pnevmatikakis and Giovannucci, 2017) and residual PMT offsets were subtracted. Residual motion artifacts or other artifacts due to instrument failure were visually identified and the corresponding recording segment excluded from further analysis. Data from mice for which strong motion artifacts occurred regularly were excluded from any analyses.

To detect ROIs corresponding to neuronal or astrocytic somata, we used different approaches adapted to the specific properties of each cell type. For astrocytes and interneuron recordings, ROIs were manually drawn around the somatic region using a previously described toolbox (https://github.com/PTRRupprecht/Drawing-ROIs-without-GUI, (Rupprecht and Friedrich, 2018)). For pyramidal cells, semi-automatic source extraction using Suite2p with subsequent manual curation of cell candidates was used (Pachitariu et al., 2019). Next, ΔF/F_0_ was computed from the fluorescence averaged across the pixels of each ROI while computing F_0_ using a running or global percentile filter that was adapted to each cell type and their respective typical timescale and activity pattern: a 15% percentile filter applied to a 67-s long running window for pyramidal cells, with a Suite2p-based neuropil subtraction with a subtraction factor of 0.15; and a 30% globally applied percentile filter for both interneurons and astrocytes.

For pyramidal cell recordings, deconvolution of the ΔF/F_0_ traces with a supervised deep network trained specifically for pyramidal cells expressing GCaMP8 (CASCADE; (Rupprecht et al., 2025, 2021)) was performed in Python to suppress shot noise and to obtain an estimate of absolute spike rates.

Cellular ROIs were manually tracked across days by careful comparison of cellular morphology and their surroundings using a custom-written tool in Matlab (Figure S2-1).

### Analysis of head-fixed behavioral recordings

The behavioral video recorded during two-photon imaging was used to extract several behavioral features. The correlation between subsequent frames of a subregion of the video (capturing forelimb and front paws) was used to compute body movement and was used as a proxy for natural, spontaneous arousal. The measured metric (1 - correlation) was scaled by the within-session maximum.

The pupil appeared bright in the behavioral camera video due to infrared laser light exiting the pupil. The equivalent diameter was computed from the area of the pupil, which was obtained from a segmentation of the pupil with standard image segmentation methods based on binarization and using the *regionprops()* function in MATLAB (MathWorks), as described before (Rupprecht et al., 2024). Occasional pupil segmentation failures were manually detected from the extracted pupil diameter traces, and then replaced by values obtained via linear interpolation.

### Quantification of cellular activity modulation by movement/arousal

To quantify the modulation of astrocyte, pyramidal cell or interneuron activity by natural arousal for two-photon imaging data, we used body movement as a proxy for natural arousal. We computed the modulation as the correlation of each cell’s activity with a time-shifted body movement for each recording segment (typically 2.5 min) within a session (typically 25 min). The time-shift (delay) was chosen to maximize the correlation of body movement with cellular activity across the entire cellular population. Delays were +3.85 s for astrocytes, +0.71 s for interneurons, and −0.28 s (almost instantaneous correlation) for pyramidal cells, in alignment with the correlation functions shown in Figure 2. To compute the same modulation index, but based on pupil diameter instead of body movement recordings, delays were +1.55 s for astrocytes, −0.64 s for interneurons, and −1.35 s for pyramidal cells. Recording segments within 60 s upon LC stimulation or shorter than 45 s were excluded from the quantification of movement modulation to prevent a confounding influence of responses to LC stimulation. The modulation indices were averaged across segments within a session by weighing according to the duration of each segment.

### Analysis of movement bout-related cellular activity

To quantify and compare cellular activity during open field-behavior and head-fixed behavior (Figure S2-6), we analyzed neural and astrocytic signals aligned to discrete movement bouts. For open-field behavior, movement bouts were identified using locomotion speed derived from the mouse’s center of mass tracked with DeepLabCut (Mathis et al., 2018). For head-fixed behavior, bouts were defined based on body motion energy as described above. Thresholds for motion detection were adjusted by visual inspection, with a single threshold applied to all open-field analyses and a single threshold applied to all head-fixed datasets. Segments shorter than 1.0 s were excluded, and movement bouts interrupted by pauses shorter than 1.0 s were merged and treated as contiguous bouts. Neural and astrocytic activity aligned to bout onset was extracted and averaged for each recording session. For improved visualization of onset and offset responses, cellular activity associated with each movement bout was temporally normalized by resampling to the median bout duration across the respective recording paradigm (3.1 s for open field behavior, 6.7 s for head-fixed behavior).

### Quantification of cellular activity modulation by LC stimulation

To quantify the modulation of astrocyte, pyramidal cell or interneuron activity by LC stimulation for two-photon imaging data, we computed the average ΔF/F in response to LC stimulation in a window following strong (≥20 Hz) LC stimulation, with the delay and duration of the window adapted to the typical responses observed for the respective cell type. The response window was chosen as the 20 s immediately after start of LC stimulation for astrocytes (to capture the slow evoked activity patterns for astrocytes), the 10-s window immediately after start of LC stimulation for interneurons (to capture both the slow inhibition and the fast activation in different interneuron subpopulations), and the 10-s window delayed by 10 s after start of LC stimulation for pyramidal cells (to capture the slow inhibition across pyramidal cells).

### Modeling of cellular activity by movement using dilated linear regression

To control for the confounding effect of movement on cellular activity, we subtracted a cross-validated linear model of cellular activity from the measured cellular activity. To model cellular activity as a function of body movement, we used a dilated variant of linear temporal regression, as previously described (Rupprecht et al., 2024). We averaged timepoints of the movement variable around the to-be-regressed timepoint into bins that exponentially increased their width with the temporal distance from the current timepoint, resulting in an effective time window of ±8.5 s (±64 timepoints sampled at 7.5 Hz for interneurons). The vector of the dilated regressor was used to linearly regress the observed activity separately for each interneuron with the glmfit() function in MATLAB. Models were fit to 80% of the data and applied to the remaining 20% to exclude overfitting. The activity modeled as a function movement was subtracted from the originally recorded activity to obtain cellular activity with tentative decontamination of the effect of movement effects. This procedure enables the improved isolation of responses specifically to LC stimulation.

### Identification of reliably responding interneurons

To identify interneurons that exhibited reliable responses towards LC stimulation and to discard instances of spurious neuronal activation, we computed a simple reliability metric. First, we retrieved for each neuron all ΔF/F responses to strong LC stimulation (≥20 Hz) within a session and extracted a time window of 100 imaging data points before to 200 imaging points after start of LC stimulation (t = −13.3 s to 26.7 s). Then, we computed the Pearson’s correlation between these peri-stimulus time traces across the repetitions of the stimulus for the same neuron within a session and computed the median correlation across pairs of repetitions as the reliability metric. For control, we repeated the same process, but with a total of 106 stimulation pairs randomly selected not from the same neuron but from different neurons. Neurons with a reliability metric exceeding a threshold (mean plus standard deviation of the random control values: >0.13) were defined as reliably responding interneurons.

### Astrocytic population vector analysis

For population vector analysis, we averaged cellular activity within a 10-s time window, resulting in a 1D vector of a length equal to the number of cells in the investigated population. We then compute the Pearson’s correlation coefficient between pairs of these 1D population vectors to obtain population vector correlation matrices as shown in Figure 3L. This analysis quantifies the similarity of pairs of population activity patterns, which, as opposed to other measures such as Euclidean distance, does not take into account the absolute magnitude of activity and only compares activity patterns.

### Sensitivity of astrocytes in response to LC stimulation

We observed that some astrocytes were already strongly activated by comparatively weaker stimulation (e.g., 5 Hz), with only a minor additional increase of responses for stronger stimulation (e.g., 20 Hz) - we term these astrocytes “more sensitive” to LC stimulation. Other astrocytes were almost not activated by weaker stimulation and required stronger stimulation to display clear responses. To compute a proxy for cell-specific “sensitivity” to LC stimulation, we therefore computed the sensitivity S5/20 as the ratio of the response to 5 Hz stimulation (Res5Hz) and the response to 20 Hz stimulation (Res20Hz):

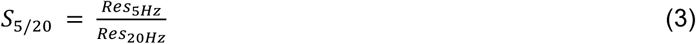

A high value of S_5/20_ means that the specific astrocyte is highly sensitive, i.e., responds already to strong responses for relatively weaker LC stimulation. A low value of S5/20 means that the specific astrocyte is less sensitive and requires strong LC stimulation to respond.

For the subset of recordings where not only 20 Hz and 5 Hz but also other stimulation strengths (1 Hz, 3 Hz, 10 Hz, 40 Hz) were additionally applied, a more complete activation function together with activation and saturation points could be extracted. We performed such tentative analyses using a sigmoid fit of the activation *A* as a function of the stimulation strength *x*, *A(x)*:

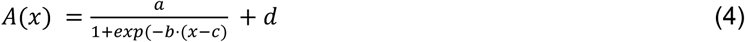

with the fit parameters *a*, *b*, *c* and *d*. Parameters *a* and *d* are scaling parameters that are less biologically relevant. Parameter *b* determines the steepness of the activation. Parameter *c*, most importantly, determines the inflection point and therefore the level of activation *x* where the astrocyte starts responding strongly to LC stimulation.

### Statistics and reproducibility

Unless otherwise indicated, nonparametric, two-sided tests were used (Mann–Whitney rank-sum test and Wilcoxon signed-rank test for unpaired and paired conditions). Box plots used standard settings in MATLAB, with the central line at the median of the distribution, the box at the 25th and 75th percentiles and the whiskers at extreme values excluding outliers.

## Supplementary Figures

**Figure 1 Supplement 1.**
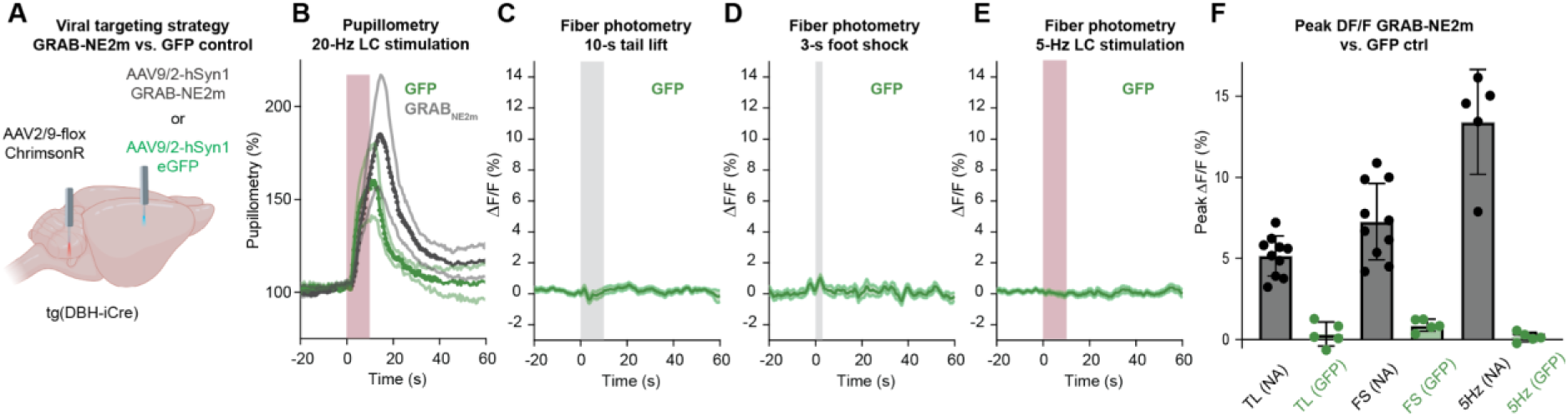
GFP control experiments related to. Figure 1**. A**, Viral targeting strategy for control (GFP) or regular (GRAB-NE2m) experiments, each performed in DBH-iCre transgenic mice together with injection of Cre-dependent ChrimsonR-tdTomato. **B**, Upon optogenetic stimulation of LC, GFP control animals (green) show a pupil response similar to GRAB-NE2m-expressing animals (gray), indicating successful targeting of LC with the optogenetic sensors (ChrimsonR) under both conditions. **C-E**, No discernible signal change in response to (**C)** tail lift, (**D)** foot shock, and (**E**) LC stimulation in GFP control animals. **F**, Comparison of peak ΔF/F of GFP control vs. GRAB-NE2m animals (GRAB-NE2m values taken from Figure 1E-G).

**Figure 1 Supplement 2.**
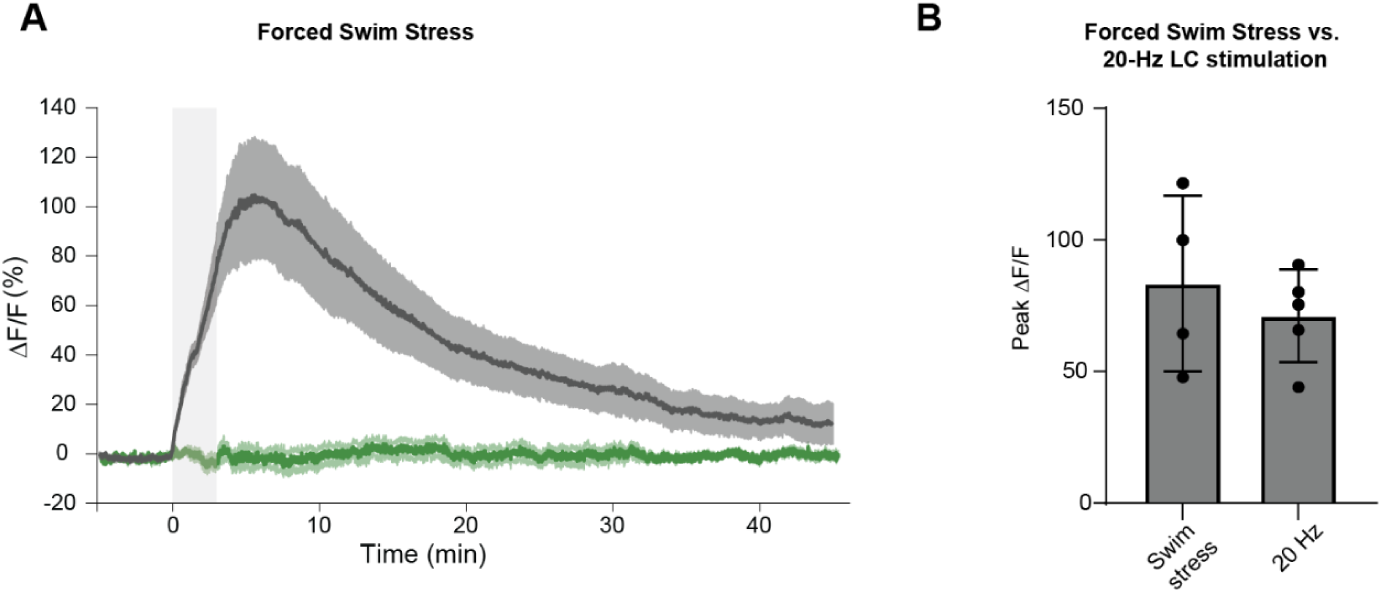
Forced swim stress experiment with photometry of NA. Mice expressing the NA sensor GRAB-NE2m were subject to a forced swim in 18°C cold water for 3 min minutes. Fiber photometry recordings were performed several minutes before, during, and after forced swim stress. **A**, Time course of fiber photometry of NA sensor-expressing mice (gray, n = 4) and control mice expressing GFP (green, n = 4). The measured NA release increases briefly even after the end of the acute stressor, and decays slowly back to baseline on a timescale of 10s of minutes. **B**, Comparison of NA release (peak ΔF/F) during forced swim stress (values from panel A) and strong optogenetic LC stimulation (values from Figure 1H). The values are of a similar magnitude, illustrating that the 20-Hz LC stimulation used in our study results in NA release similar to a strong stressor. Due to tightening regulations on animal welfare, the swim stress paradigm was not included for subsequent experiments with astrocytic and neuronal fiber photometry.

**Figure 1 Supplement 3.**
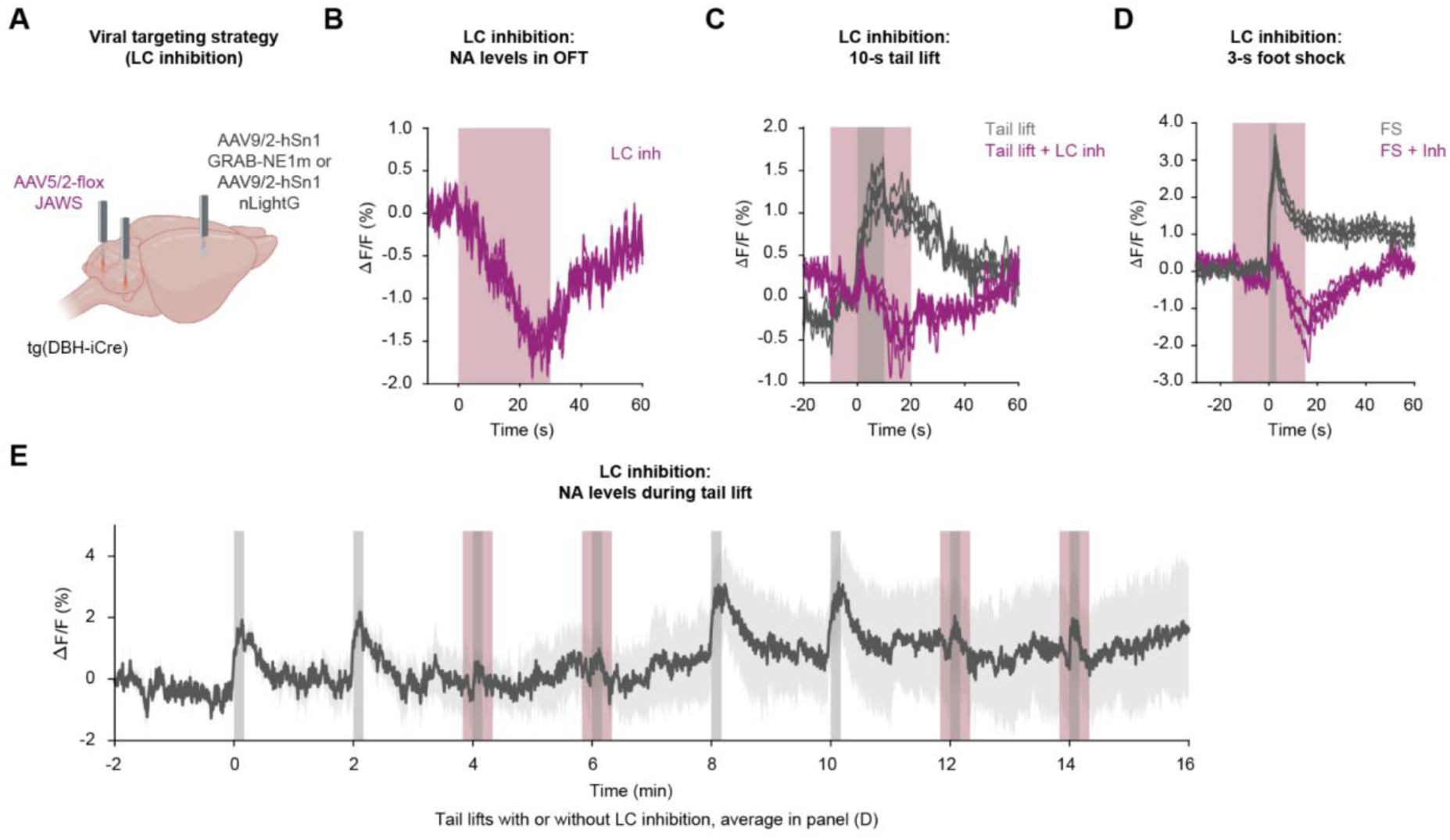
Optogenetic inhibition of LC prevents NA release during stress. **A**, Viral targeting strategy to inhibit LC activity (bilateral injection of Cre-dependent Jaws in DBH-iCre mice; bilateral fiber implants) while monitoring NA levels (NA sensors GRAB-NE1m or nLightG) in HC. Inhibition of LC with Jaws decreases NA levels (**B**) at baseline, (**C**) during tail lift, and (**D**) during foot shock (without LC inhibition in green; with LC inhibition in purple). **E**, Example fiber photometry recording with responses to tail lift with (gray+red shading) and without inhibition (gray shading only).

**Figure 1 Supplement 4.**
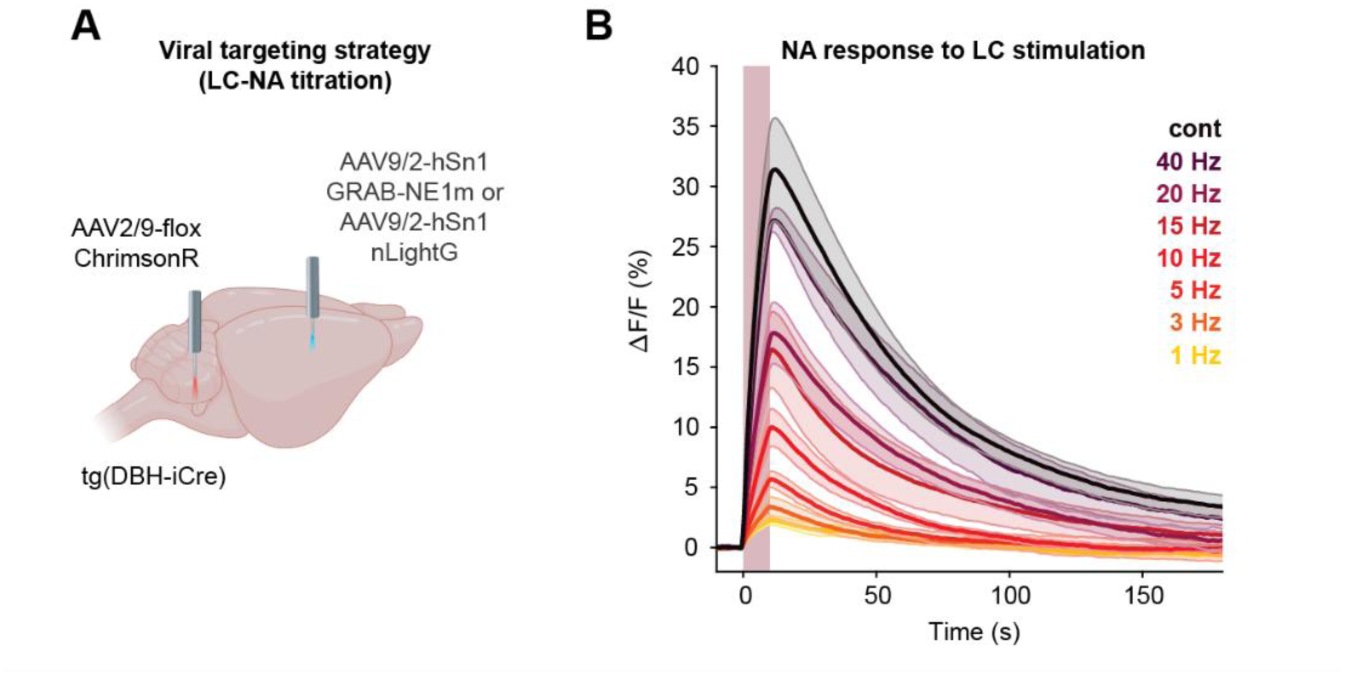
NA release in HC upon optogenetic LC stimulation. **A**, Viral targeting strategy to optogenetically activate LC (injection of Cre-dependent ChrimsonR in DBH-iCre mice) while monitoring NA levels (NA sensors GRAB-NE1m or nLightG) in HC. **B**, NA release in response to increasing LC stimulation intensities (10 s of 1 Hz, 3 Hz, 5 Hz, 10 Hz, 15 Hz, 20 Hz, 40 Hz and continuous stimulation, 10-ms pulses, 10 mW laser power, for n = 4 mice).

**Figure 2 Supplement 1.**
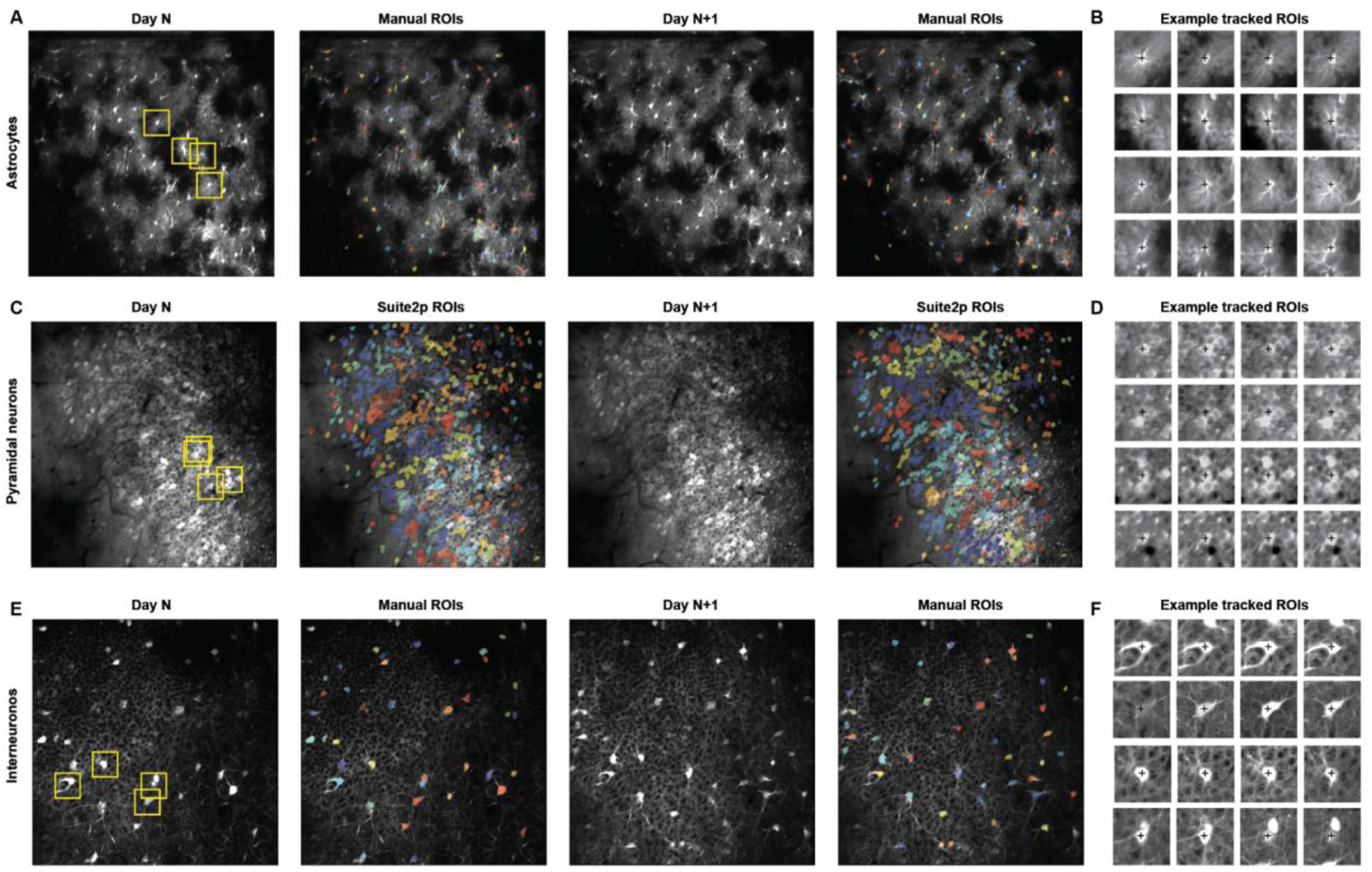
Tracking of astrocytes, pyramidal cells and interneurons across days. **A**, An astrocytic FOV (central plane of three imaging planes) with manually selected somatic ROIs (colored overlays) for two separate days. The coloring of ROIs does not reflect cell identity across days. **B**, Example ROIs covering the same astrocytes tracked across days. ROI locations are highlighted with yellow squares in (A). Across 4 mice, 960 cells were tracked in ≥2 sessions, 646 in ≥3 sessions, and 406 in ≥4 sessions. **C**, A pyramidal cell FOV with sources of neuronal activity, extracted with Suite2p (colored overlays), for two separate days. **D**, Example ROIs covering the same neurons tracked across days. ROI locations are highlighted with yellow squares in (C). Across 3 mice, 897 pyramidal cells were tracked in ≥2 sessions, 619 in ≥3 sessions, and 324 in ≥4 sessions. **E**, An interneuron FOV (second plane out of four planes, counted from *stratum radiatum* to *stratum oriens*) with manually selected somatic ROIs (colored overlays) for two separate days. **F**, Example ROIs covering the same cells tracked across days. ROI locations are highlighted with yellow squares in (E). Across 4 mice, 553 interneurons were tracked in ≥2 sessions, 481 in ≥3 sessions, and 385 in ≥4 sessions.

**Figure 2 Supplement 2.**
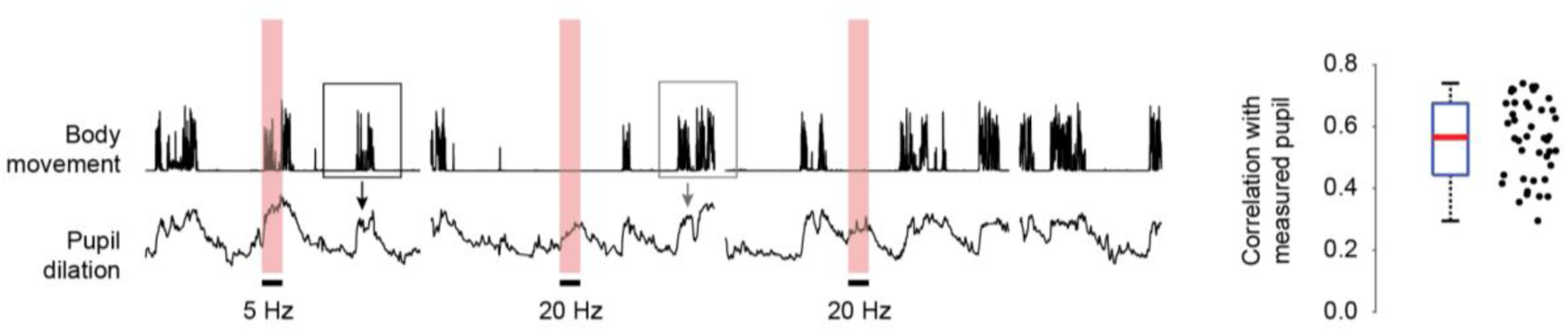
Body movement as a proxy for endogenous arousal. Visually, pupil dilation appeared as a lowpass-filtered version of body movement. Importantly, as described previously (Rupprecht et al., 2024), body movement reflected not only locomotion but also locomotion-independent body movement that was reflected by pupil dilation. To demonstrate that body movement is predictive of pupil dilation (and therefore arousal), we used dilated linear regression (see Methods) to predict pupil dilation from a temporally extended window of body movement (left). We found that the predicted pupil dilation correlated highly with the experimentally measured pupil diameter (median correlation ± S.D., 0.57 ± 0.13 across 42 imaging sessions from 13 animals). Lower values within this distribution are likely due to imprecisions of pupil diameter measurement because of unstable lighting conditions (e.g., we used an UV LED pointed at the mouse’s eye to preconstrict the pupil; eye movements of the mouse towards or away from the UV light therefore result in pupil dilations or constrictions) and subsequent pupil segmentation errors, as well as from pupil dilations not stemming from natural arousal but from optogenetic LC stimulation (indicated in red here).

**Figure 2 Supplement 3.**
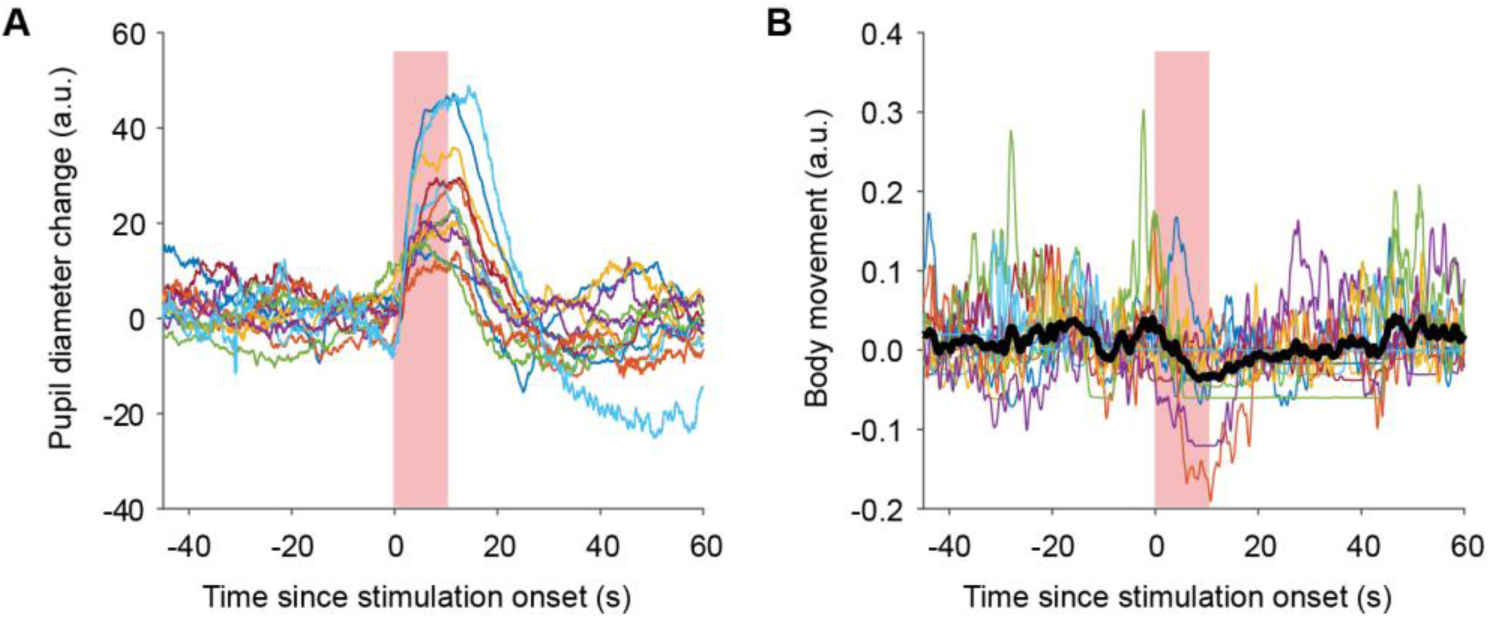
Behavioral responses upon LC stimulation for two-photon microscopy experiments. **A**, Pupil diameter changes upon stimulation with 10-s 20-Hz protocols were recorded and averaged across stimulations for each imaging session and then averaged across sessions for each animal. For each animal (n = 13), the pupil diameter change is shown with a different color. While the magnitude of evoked pupil diameter changes is variable across animals, each of them shows a clear pupil diameter increase upon stimulation, indicating successful optogenetic stimulation of LC. **B**, Body movement upon stimulation with 10-s 20-Hz protocols as in (A) (n = 13 animals). On average across animals (blue trace), animals exhibit a reduction of body movement, consistent with previous reports of behavioral arrest upon strong and long-lasting LC stimulation. One animal consistently exhibited brief motion bouts upon LC stimulation (medium blue trace; the neuronal activity extracted from this mouse was therefore analyzed separately from all other mice in Fig. S5-1).

**Figure 2 Supplement 4.**
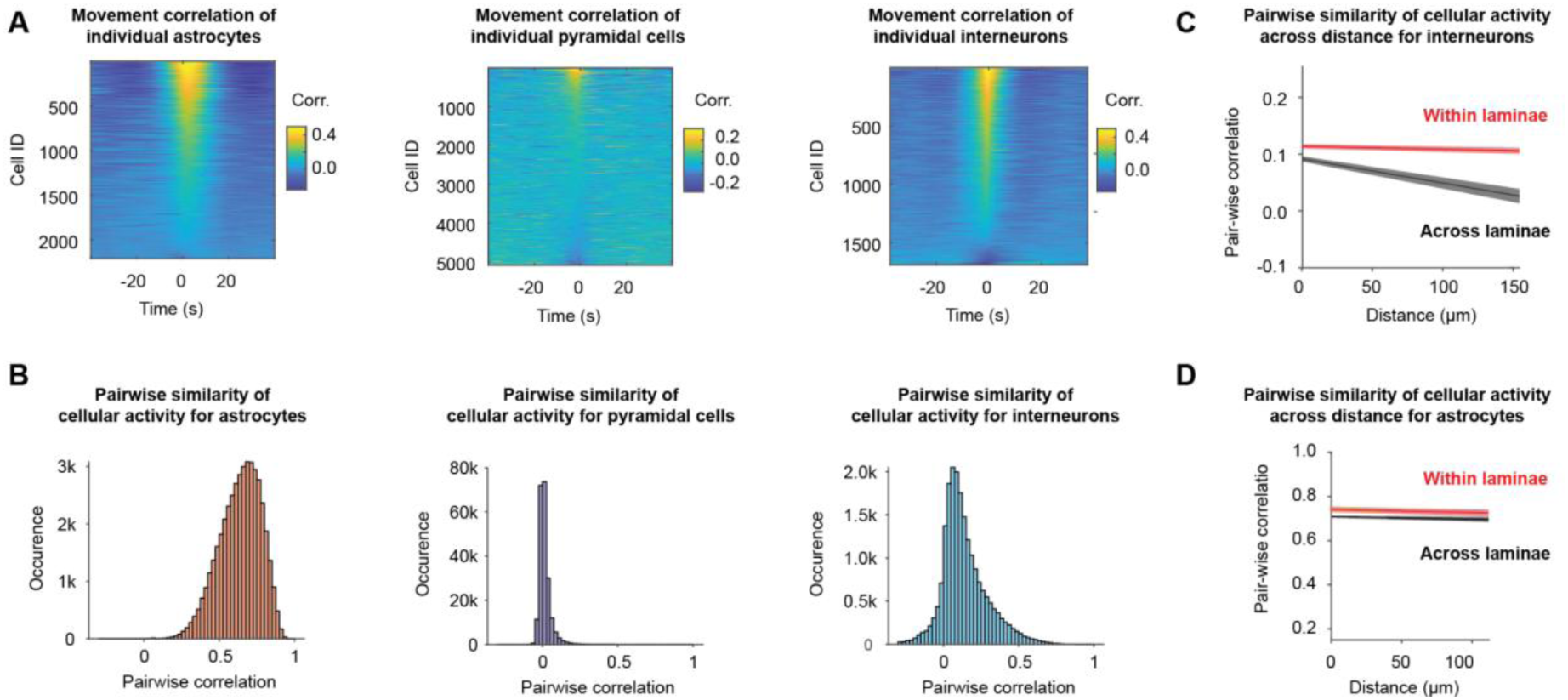
Single-cell correlation functions and pairwise correlations for all cell types. **A**, Correlation function of cellular activity with body movement as a proxy for arousal as shown in Figure 2F,J,N, but for individual cells. The cells are sorted according to the value of the movement modulation index of each cell (see Methods). **B**, Pairwise correlation between the activity patterns of all pairs of simultaneously recorded cells. **C**, Pairwise correlation as in panel (B) but plotted as distance-dependent fit to the data (fit value with 95% confidence interval of the linear fit as shading). For the condition “within laminae”, cell pairs within an imaging plane are considered, presumably within very close laminar distance. For the condition “across laminae”, cell pairs from non-identical imaging planes are considered only. The result demonstrates that interneuron activity is relatively similar within but more different across laminae. **D**, Same as in panel (C) but for astrocytes. The result demonstrates that astrocyte activity is similarly synchronized within as well as across laminae.

**Figure 2 Supplement 5.**
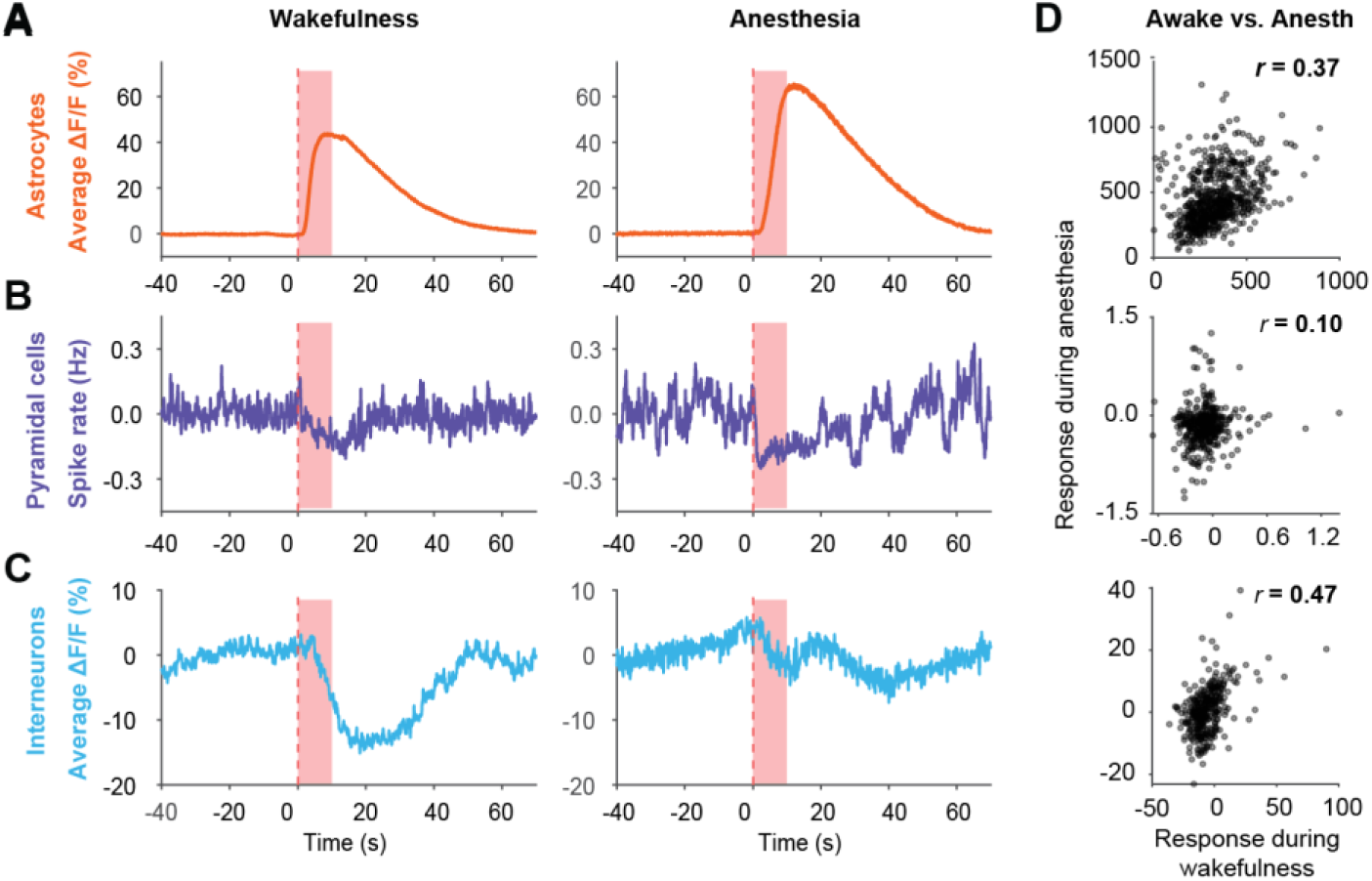
Cell-type specific responses to LC stimulation during anesthesia with two-photon microscopy. Average responses (average across all cells, animals and sessions) to LC stimulation for astrocytes (**A**), pyramidal cells (**B**) and interneurons (**C**) during wakefulness (left) and anesthesia (right). Under both conditions, similar overall responses can be observed (activation for astrocytes, inhibition for pyramidal cells and interneurons). For interneurons, a slow activation not observed during wakefulness can be seen riding on top of the inhibition. **D**, Responses on the cellular level (responses to LC stimulation computed as in Figures 3 to 5) are correlated across states for astrocytes and interneurons but not for pyramidal cells, consistent with results observed for response stability across sessions during wakefulness (Figures 3 to 5).

**Figure 2 Supplement 6.**
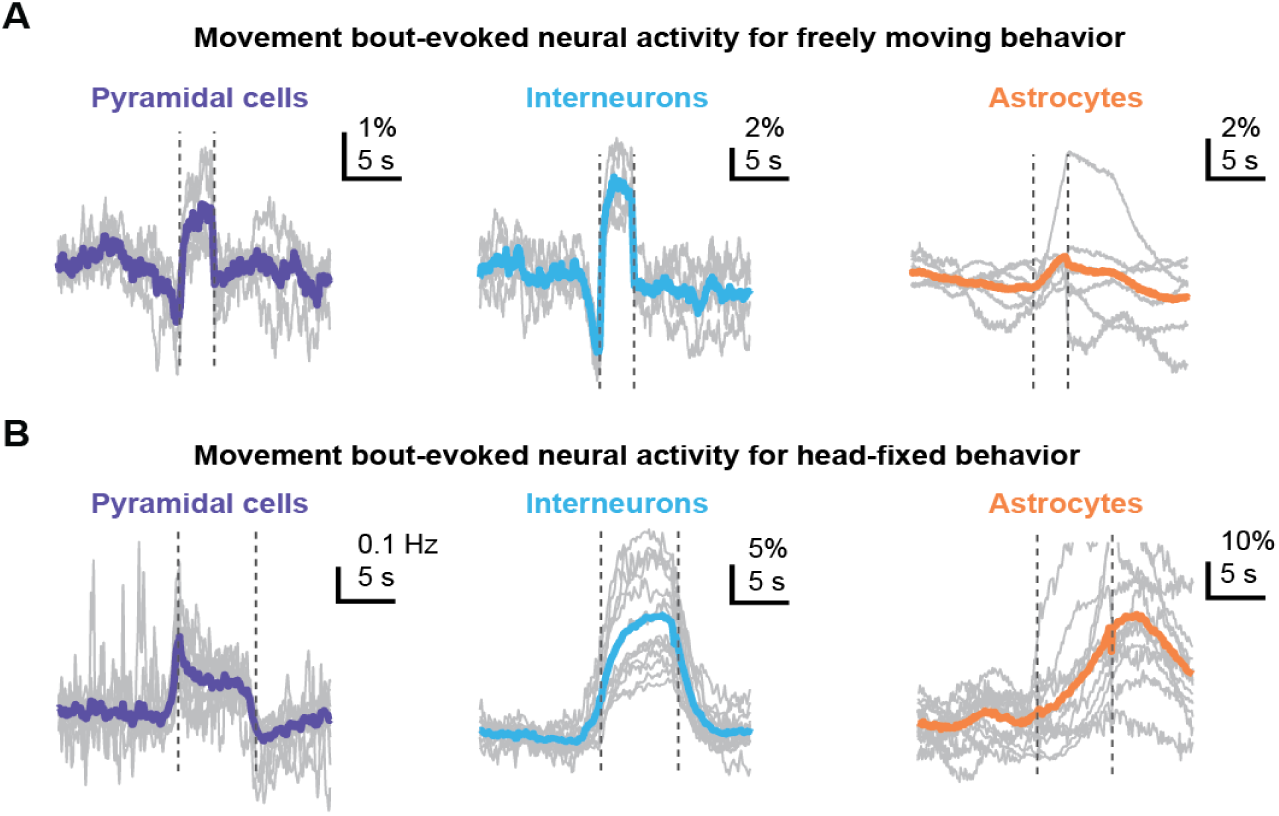
Cell-type specific modulation during freely moving vs. head-fixed behavior. To compare movement bout-evoked cellular activity between fiber photometry experiments in freely moving mice (Figure 1) and two-photon imaging experiments in head-fixed mice (Figure 2), we applied the same criteria to detect contiguous movement bouts (Methods). For each behavioral paradigm, the cellular response to each bout was temporally normalized by resampling to the median bout duration (3.1 s for freely moving behavior; 6.7 s for head-fixed behavior). Averages were performed across bouts within a session (gray traces) and then averaged across sessions to obtain the grand average (colored traces) (freely moving: 6 sessions from 6 mice for astrocytes, 5 sessions from 5 mice for interneurons, 5 sessions from 5 mice for pyramidal cells; head-fixed: 12 sessions from 4 mice for astrocytes, 14 sessions from 4 mice for interneurons, 12 sessions from 5 mice for pyramidal cells). **A**, Movement-evoked activity during freely moving behavior. **B**, Movement-evoked activity during head-fixed behavior. The longer median bout durations and the stronger astrocytic response observed during head-fixed behavior may reflect the higher effort necessary to initiate locomotion and the presumably more stressful mismatch between intended and actual movement under head-fixed conditions. These observations suggest that movement during head-fixed behavior adds a stressful component that is less prominent during freely moving behavior in an open field.

**Figure 3 Supplement 1.**
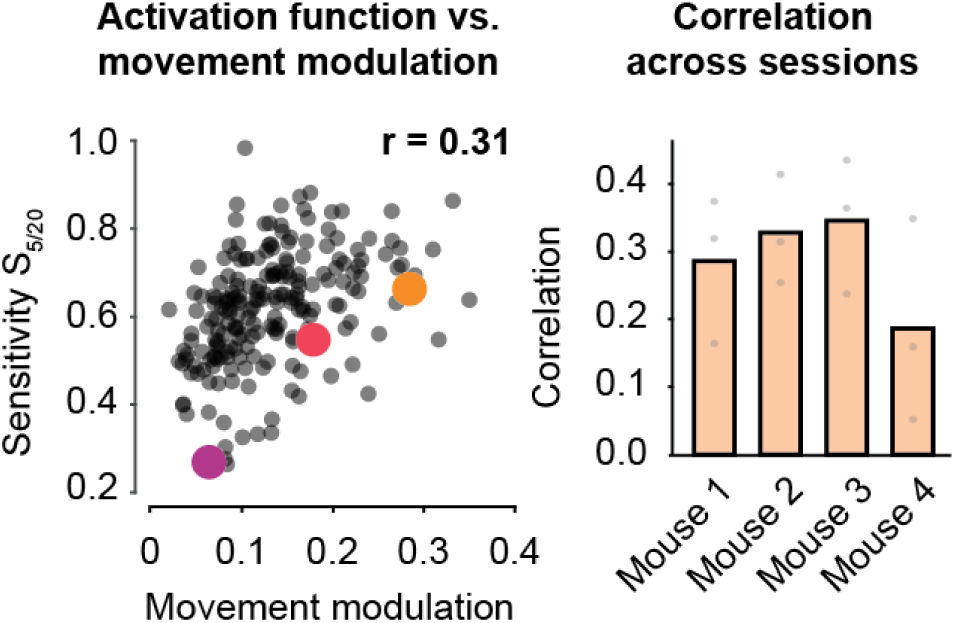
Correlation between sensitivity to LC responses and response to movement. *Left:* Correlation of responses to LC stimulation sensitivity as described in Figure 3I and movement modulation for an example session. Individual dots refer to individual neurons. *Right:* All Pearson correlation coefficients for 12 sessions from 4 animals.

**Figure 4 Supplement 1.**
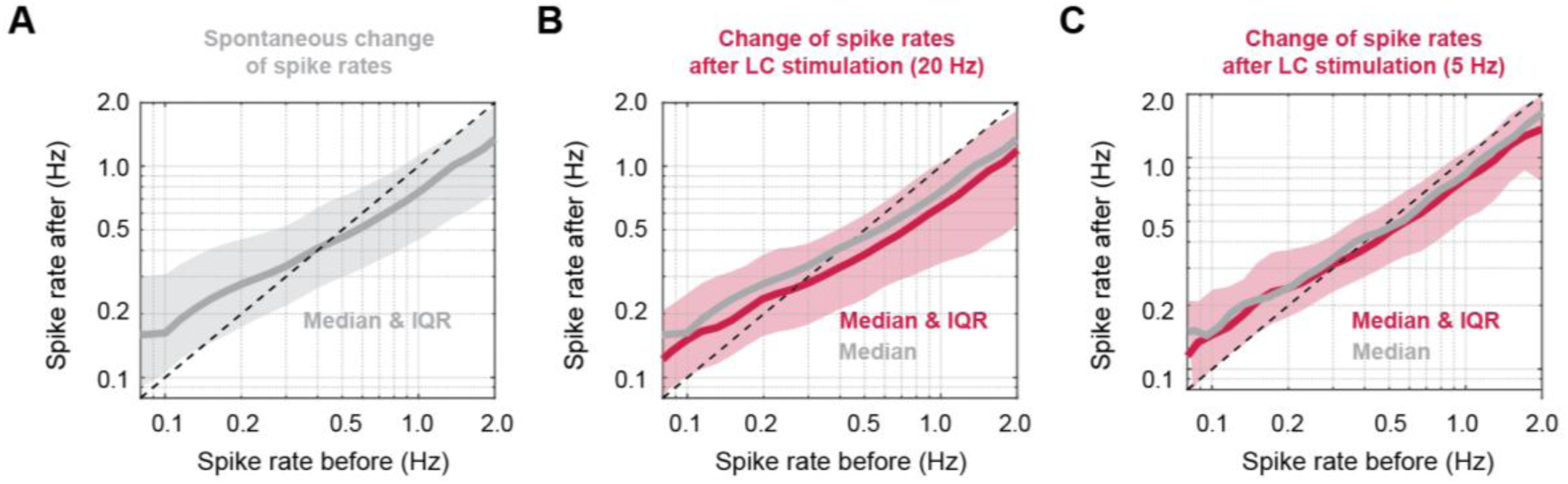
LC-induced change of spike rates, related to. Figure 4D. Change of spike rate from a 10-s pre-stimulus window (x-axis) to the 10-s window of LC stimulation (y-axis) as in Figure 4D. **A**, Same as in Figure 4D but only for spontaneous transitions (no LC stimulation), shown as median of the distribution together with the inter-quartile range as shaded corridor. **B**, Same as in Figure 4D but only for transitions induced by LC stimulation, shown as median of the distribution (red) together with the inter-quartile range as a shaded corridor. For reference, the median of the distribution for spontaneous transitions is shown in gray. **C**, Same as in panel (B) but for 5-Hz instead of 20-Hz LC stimulation. As in Figure 4D and panels (A,B), an - albeit smaller - reduction of the spike rate upon LC stimulation can be observed across all pre-stimulation spike rates..

**Figure 5 Supplement 1.**
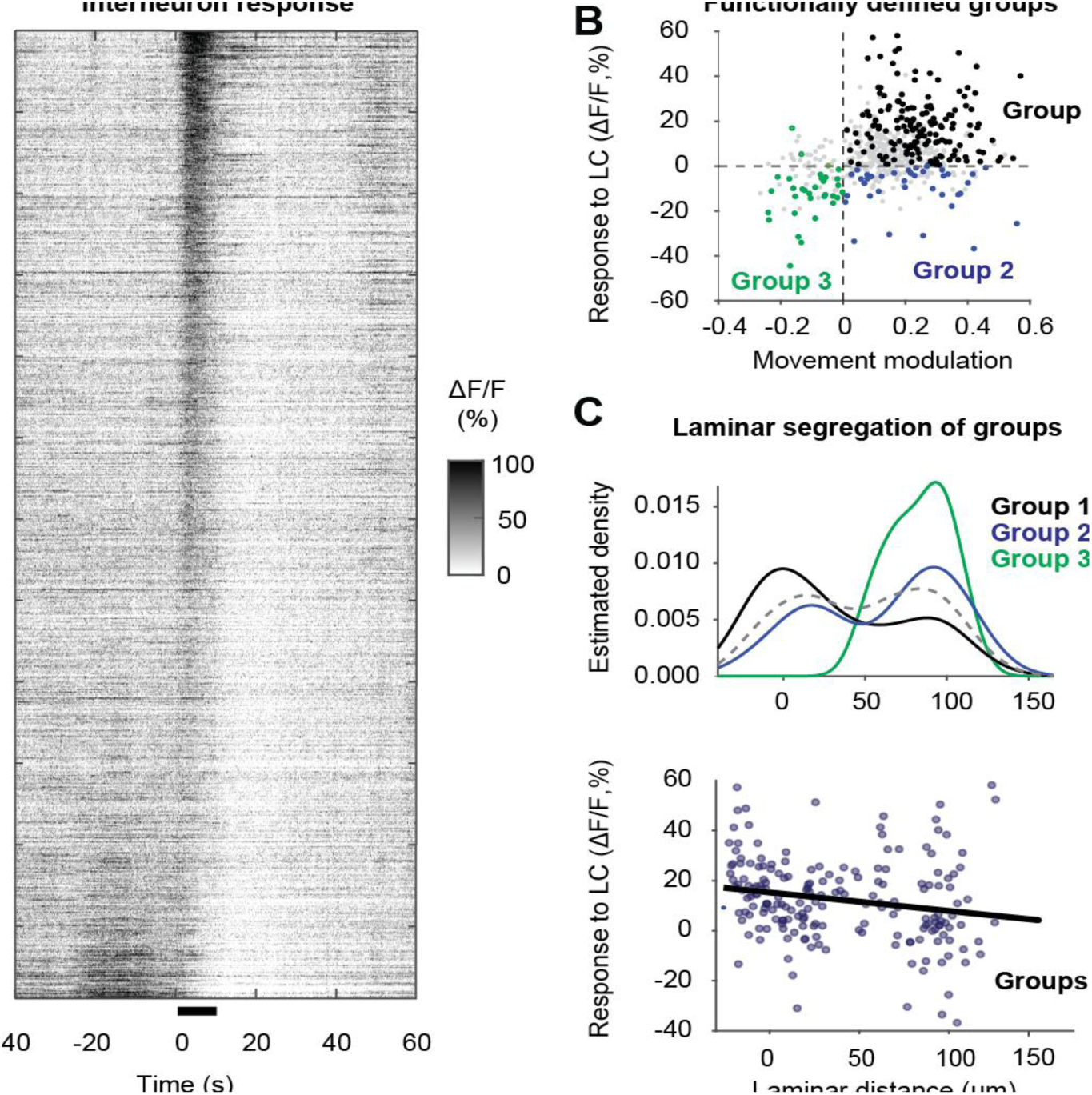
Additional analyses related to. Figure 5 **(outlier mouse).** Additional analyses for the outlier mouse, which was excluded from the analyses in Figure 5 due to LC stimulation-triggered movement bouts. Despite this potential confound, all analyses performed for this mouse only reflected the effects observed for the other mice in Figure 5. **A**, Interneuron responses to 20 Hz LC stimulation, sorted by the response during the 10-s stimulation window. Data pooled from 3 sessions. This mouse exhibits higher apparent responses to LC stimulation compared to the data shown in Figure 5A, but this difference may be explained by movement systematically triggered by LC stimulation in this specific mouse. **B**, Relationship of natural arousal (x-axis, quantified by body movement modulation of each cell) and response to LC stimulation (y-axis), analysis performed as for Figure 5G, resulting in three primary groups: Group 1, which responds positively to both LC stimulation and movement (black); Group 2, which responds positively to movement but is primarily inhibited by LC stimulation (blue); and Group 3, which is inhibited both by LC stimulation and negatively modulated by body movement (red). **C**, *Top:* Laminar distribution of the three cell groups shown in panel (B), analogous to Figure 5H. “Zero” on the x-axis indicates the position of the pyramidal cell layer, with positive distance going deeper into the stratum oriens. Neurons responsive to LC stimulation (black, Group 1) are located closer to the pyramidal cell layer. Neurons negatively modulated by movement and LC stimulation (red, Group 3) are primarily located in deep layers close to the corpus callosum. Neurons from Group 2 are distributed across all recorded laminae. The overall distribution of recorded interneurons is shown in the background as a gray dashed line. *Bottom:* Neurons responding positively to movement (Groups 1 and 2) increase their response strength to LC stimulation when located more superficially, i.e., closer to the pyramidal cell layer, as shown for the other mice in Figure 5I.

**Figure 5 Supplement 2.**
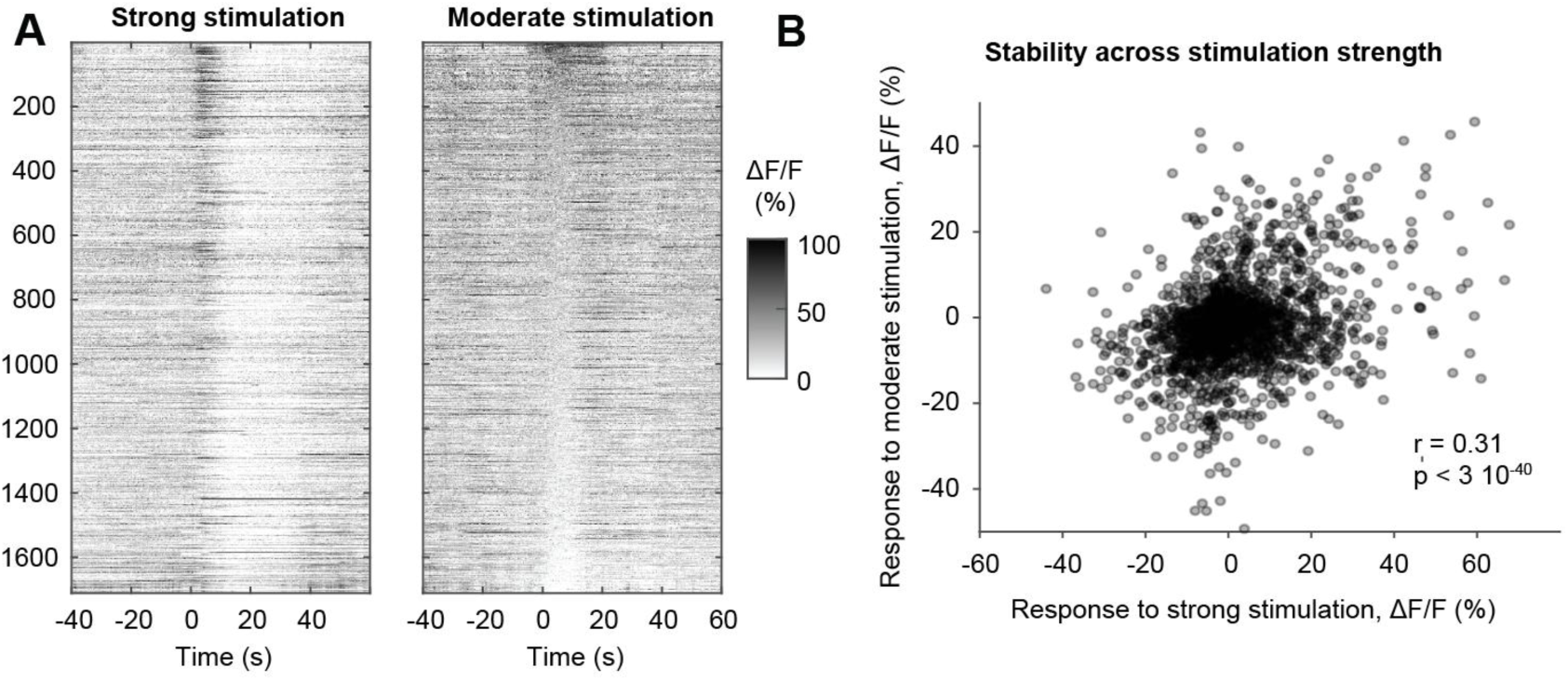
Stability of interneuron responses to moderate (5 Hz) vs. strong (≥20 Hz) LC stimulation. A, Interneuron responses to ≥20-Hz LC stimulation (left) and 5-Hz LC stimulation (right), sorted by the response during the 10-s stimulation window for 5-Hz stimulation. The sorted suggests the neurons that are inhibited/activated by strong stimulations of LC are also inhibited/activated by moderate stimulations of LC. Data pooled from all animals (N = 4) across all recorded cells (n = 1710 neurons). **B,** Correlation of responses to strong vs. moderate stimulation strength, measured as the response magnitude during the 10-s stimulation window, with the statistical outcomes of the computed correlation across all neurons.

**Figure 5 Supplement 3.**
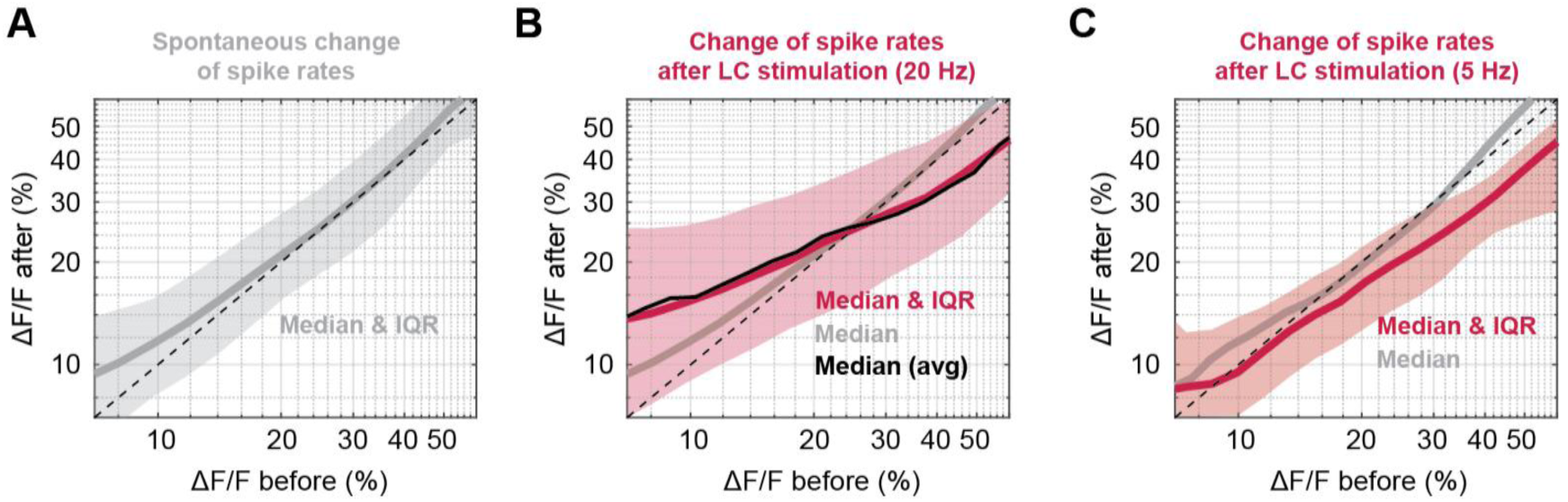
LC-induced change of ΔF/F, related to. Figure 5D. Change of ΔF/F from a 10-s pre-stimulus window (x-axis) to the 10-s window of LC stimulation (y-axis) as in Figure 5D. **A**, Same as in Figure 5D but only for spontaneous transitions (no LC stimulation), shown as median of the distribution together with the inter-quartile range as shaded corridor. **B**, Same as in Figure 5D but only for transitions induced by LC stimulation, shown as median of the distribution (red) together with the inter-quartile range as a shaded corridor. For reference, the median of the distribution for spontaneous transitions is shown in gray. In addition, we permuted stimulation trials across repetitions for individual neurons and performed the same analysis (black), resulting in no obvious difference compared to the experimental data (red). This control analysis indicates that interneurons do not show a specific sensitivity to LC stimulation based on their activity prior to stimulation, but rather a specific sensitivity to LC stimulation that is coupled with a specific baseline activity level: interneurons with lower baseline activity are more likely to be activated by LC stimulation, while interneurons with higher baseline activity are more likely to be inactivated by LC stimulation. **C**, Same as in panel (B) but for 5-Hz instead of 20-Hz LC stimulation. For this lower stimulation intensity, the inactivation of highly active interneurons can still be observed, while the activation of inactive neurons as observed for 20-Hz stimulation is not present.

**Figure 5 Supplement 4.**
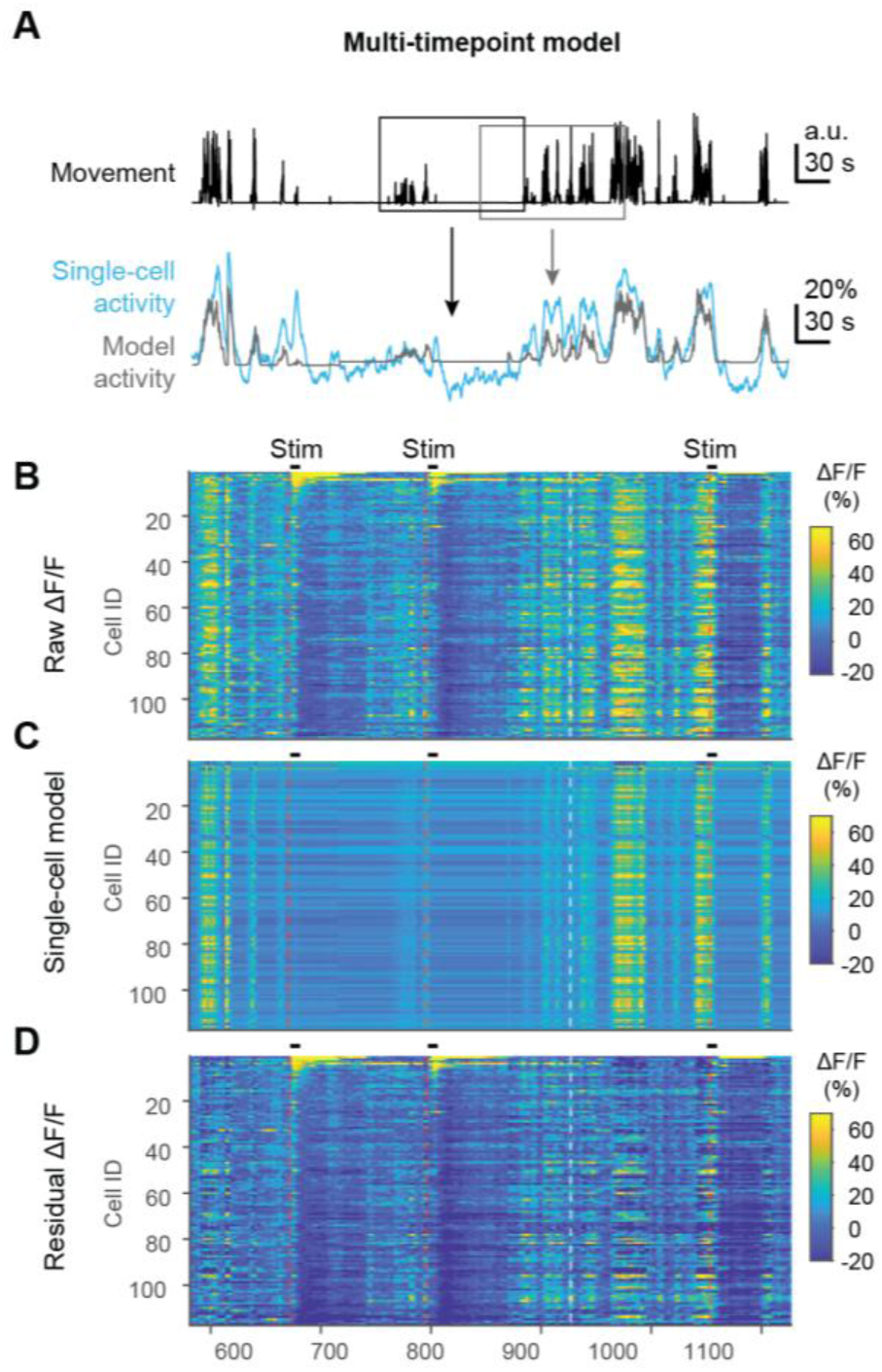
Multi-timepoint modeling to explain interneuron activity with movement. The modeled activity patterns were subtracted from experimentally recorded data, resulting in a residual pattern that is not or less contaminated by the influence of body movement, thereby providing a cleaner readout of responses to LC stimulation. **A**, A model based on dilated linear regression (see Methods) was trained in a cross-validated manner for each neuron to explain its activity (ΔF/F) as a linear function of current and past body movement. Movement (top) is used as the regressor to explain single-cell activity (bottom, blue) with a linear model (bottom, red). **B**, Example of raw imaging data (ΔF/F), excerpt from a single recording session. **C**, Single-neuron model of the activity patterns seen in (B), based on body movement as regressor. **D**, Residual activity patterns, obtained by subtracting the modeled activity pattern (C) from the original activity pattern (D). It is clear that the effect of body movement (compare with panel A) is reduced, although not completely eliminated.

**Figure 5 Supplement 5.**
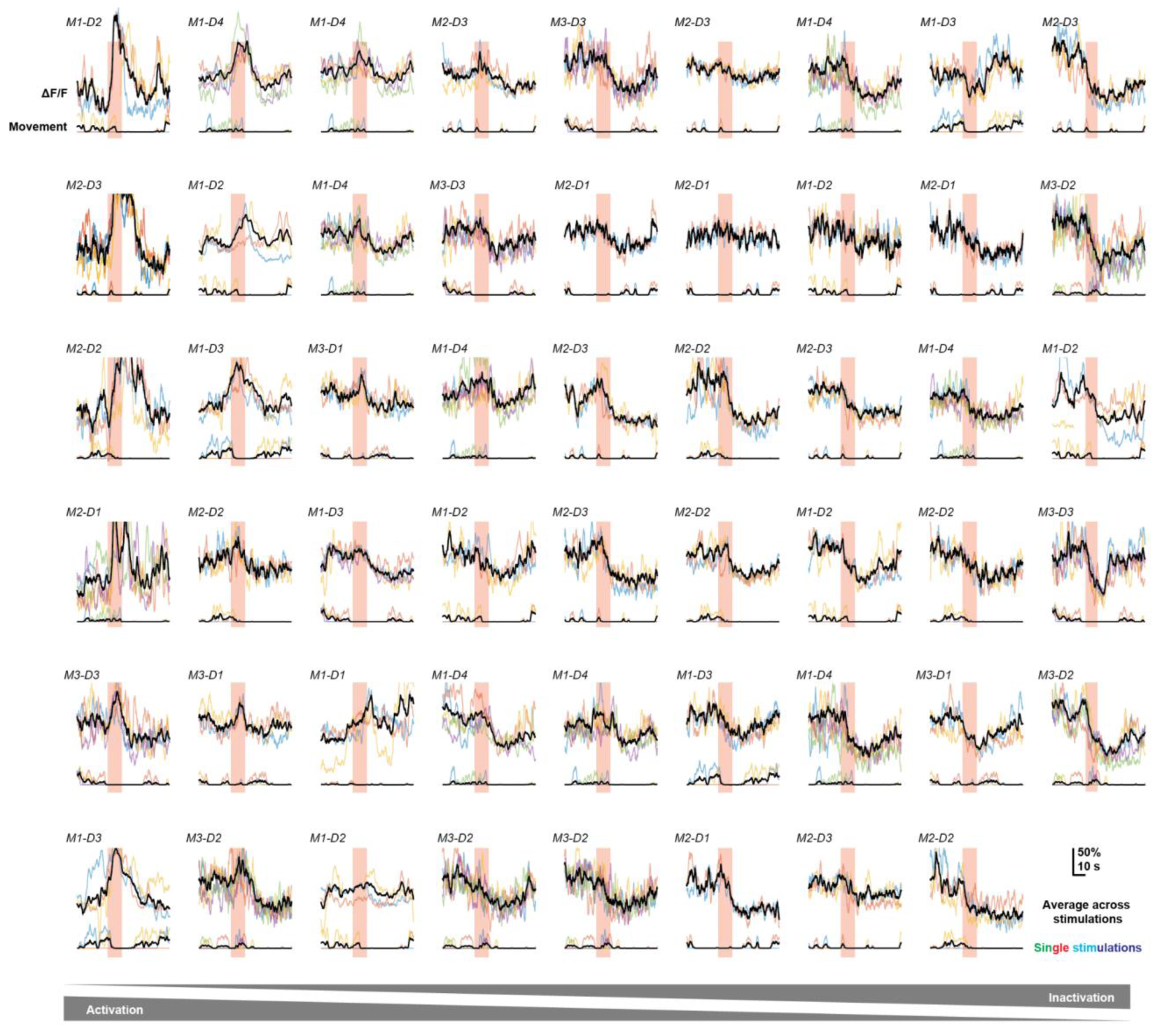
Example responses of individual interneurons to LC stimulation (≥20 Hz). Only neurons that exhibited reliable responses (see Methods) are shown. They are sorted according to the response magnitude during the 10-s stimulation window in a columnar sequence (neurons shown in the left-most column show strongest activating response; neurons in top row of this column show the strongest activating response within this column). Individual stimulation instances are shown with coloring, average across responses in black. Traces of body movement as a proxy for natural arousal and potential confound are shown below, with the colors matching between corresponding trials. Subset of neurons with reliable responses (every 8th neuron) for mice 1-3, upon which most analyses in Figure 5 are based.

**Figure 5 Supplement 6.**
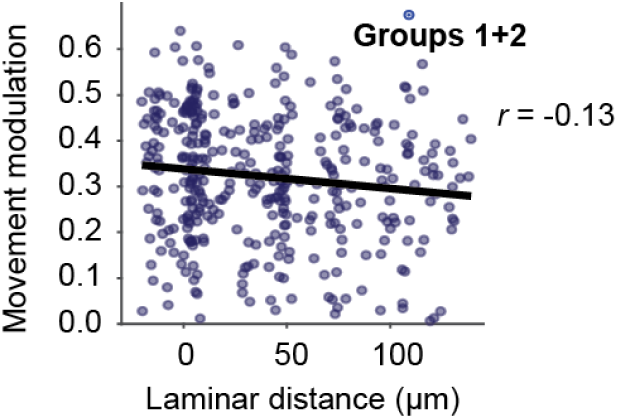
Movement modulations as a function of laminar location within deep CA1. Neuronal responses to movement of Group 1 and 2 interneurons are not well explained by laminar depth (r = −0.13). This is in contrast to responses to LC stimulation, which are well explained by laminar depth for the same set of interneurons (Figure 5J).

**Figure 6 Supplement 1.**
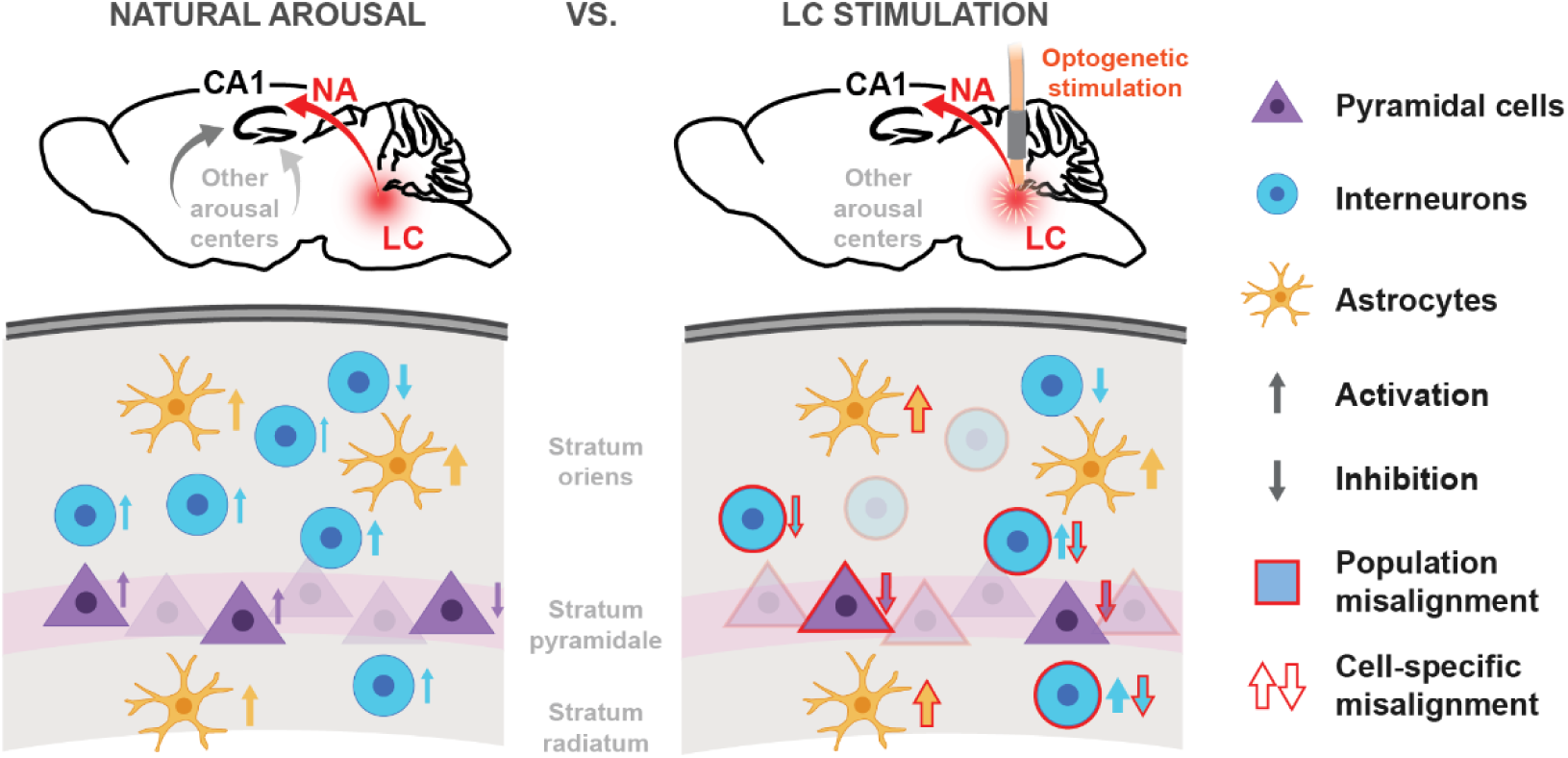
Schematic overview of cell-type-specific effects of natural arousal and LC stimulation. Individual pyramidal cells (purple triangles) are consistently activated (upward-pointing arrows) or inhibited (downward-pointing arrows) during arousal, whereas LC stimulation induces an non-specific, slow inhibitory response. Similarly, individual interneurons (turquoise circles) are consistently activated or inhibited during arousal, but also exhibit consistent responses to LC stimulation, with a transient activation of interneurons located near *stratum pyramidale* on top of a broader slow inhibitory response. In contrast, astrocytes (orange symbols) are activated during arousal and LC stimulation. At the level of individual cells, responses to LC stimulation differ from arousal responses for all three cell types (red outlines of arrows). At the population level, pyramidal cells and interneurons differ markedly between arousal and LC stimulation (red outlines of cell symbols), while astrocyte population responses are aligned across conditions.

## Supplementary Video Legends

**Supplementary Video 1 - Calcium imaging of astrocytes in hippocampal CA1 during head-fixed behavior.**

Example of simultaneous recording of single-cell astrocytic calcium signals using multi-plane two-photon calcium imaging, together with monitoring of head-fixed behavior. Initially, the video shows a single imaging plane (top) and an overview of the behavioral monitoring camera (bottom). Then, all three simultaneously imaged planes are shown, with the top-left plane as the ventral-most and the bottom-left as the dorsal-most imaging planes. Next, example cells are selected and their extracted fluorescence traces together with behavioral variables (body movement; pupil diameter) are shown. LC stimulations are indicated as red overlaid vertical bars.

**Supplementary Video 2 - Calcium imaging of pyramidal cells in hippocampal CA1 during head-fixed behavior.**

Example of simultaneous recording of single-cell calcium signals of pyramidal cells using single-plane two-photon calcium imaging, together with monitoring of head-fixed behavior. Initially, the video shows the imaging plane (top) and an overview of the behavioral monitoring camera (bottom). Then, example cells are selected and their extracted fluorescence traces together with inferred spike rates and behavioral variables (body movement; pupil diameter) are shown. LC stimulations are indicated as red overlaid vertical bars.

**Supplementary Video 3 - Calcium imaging of interneurons in hippocampal CA1 during head-fixed behavior.**

Example of simultaneous recording of single-cell calcium signals of interneurons using multi-plane two-photon calcium imaging, together with monitoring of head-fixed behavior. Initially, the video shows a single imaging plane (top) and an overview of the behavioral monitoring camera (bottom). Then, all four simultaneously imaged planes are shown, with the top-left plane as the ventral-most and the bottom-right as the dorsal-most imaging planes. Next, example cells are selected and their extracted fluorescence traces together with behavioral variables (body movement; pupil diameter) are shown. LC stimulations are indicated as red overlaid vertical bars.

